# The Siglec-sialic acid-axis is a target for innate immunotherapy of glioblastoma

**DOI:** 10.1101/2022.11.07.515406

**Authors:** Philip Schmassmann, Julien Roux, Alicia Buck, Nazanin Tatari, Sabrina Hogan, Jinyu Wang, Sohyon Lee, Berend Snijder, Tomás A. Martins, Marie-Françoise Ritz, Tala Shekarian, Deniz Kaymak, Marta McDaid, Michael Weller, Tobias Weiss, Heinz Läubli, Gregor Hutter

**Affiliations:** Brain Tumor Immunotherapy Lab, Department of Biomedicine, University Hospital and University of Basel, Switzerland; Bioinformatics Core Facility, Department of Biomedicine, University Hospital and University of Basel, Switzerland; Swiss Institute of Bioinformatics, Basel, Switzerland; Department of Neurology, Clinical Neuroscience Center, University Hospital and University of Zurich, Switzerland; Cancer Immunotherapy Lab, Department of Biomedicine, University Hospital and University of Basel, Switzerland; Institute of Molecular Systems Biology, ETH Zurich, Switzerland; Division of Oncology, Department of Theragnostics, University Hospital Basel, Switzerland; Department of Neurosurgery, University Hospital Basel, Switzerland

## Abstract

Glioblastoma (GBM) is the most aggressive form of primary brain tumor, for which effective therapies are urgently needed. Cancer cells are capable of evading clearance by phagocytes such as microglia and monocyte-derived cells through engaging tolerogenic programs. Here, we found that high level of Siglec-9 expression correlates with reduced survival in GBM patients. Using conditional knockouts of Siglec-E, the murine functional homologue of Siglec-9, together with single-cell RNA sequencing, we demonstrated significant pro-phagocytosis effects in microglia and monocyte-derived cells in the absence of Siglec-E. Loss of Siglec-E on monocyte-derived cells enhances antigen cross-presentation and production of pro-inflammatory cytokines, resulting in more efficient T cell priming. This bridging of innate and adaptive responses delays tumor growth and results in prolonged survival. Further, we showed synergistic activity of Siglec-E blockade in combinatorial immunotherapies and demonstrate its translational potential against GBM.

## Introduction

Glioblastoma (GBM) is a fatal disease without effective long-term treatment options. The current standard of care consists of tumor resection followed by adjuvant chemoradiotherapy resulting in a median overall survival of only 14 months [1]. Cancer immunotherapy using immune checkpoint inhibitors (ICI), has improved the outcomes of different cancer patients [2], but clinical trials of systemic T cell ICI showed only disappointing results in GBM [3–5]. This was attributed in part to the highly immunosuppressive immune tumor microenvironment (iTME) of GBM, which mainly consists of yolk sac-derived microglia (MG) and monocyte-derived cells (MdCs) [6, 7], together termed glioma-associated microglia/macrophages (GAMs). Recent work identified GBM associated MG and MdCs as effector cells of tumor cell phagocytosis in response to blockade of the ‘don’t eat me’-signal CD47 [8–10]. However, variability in the magnitude and durability of this response suggests the presence of additional, yet unknown such signals.

Upregulation of sialic acid-containing glycans on tumor cell surface and in the tumor microenvironment (hypersialylation) is a key change in malignant tissue and capable of impacting tumorigenesis by promoting cell invasion and metastatic potential [11–14]. By engaging immunomodulatory sialic acid-binding immunoglobulin-like lectins (Siglecs), tumor hypersialylation can trigger tolerogenic programs in different immune cell types and contributes to the establishment of the immunosuppressive iTME [15]. Recent work has shown that inhibitory CD33-related Siglecs, including human Siglec-7, Siglec-9 and Siglec-10, promote tumor progression in various models of pancreatic, breast and ovarian cancer by inducing a regulatory M2-like phenotype in tumor-associated macrophages (TAMs) [15–18]. Similarly, increased density of sialylated glycans on cancer cells inhibits human NK cell activation and cytotoxicity [19] and facilitates induction of regulatory T cells (Tregs) through engagement of Siglec-7/-9 [20, 21]. However, little is known on the induction of tolerogenic programs via Siglec receptors on MG and MdCs in the GBM iTME.

Here, we aimed to define the role of inhibitory Siglecs in innate-centered GBM immunotherapy. We found high *SIGLEC9* expression to be associated with worse clinical outcome in glioma patients, and identified Siglec-E, the murine functional homologue of Siglec-9 [18], as an anti-phagocytic signal in a preclinical GBM model. Further we showed synergistic activity of Siglec-E blockade in combinatorial immunotherapies and demonstrated its translational potential against GBM.

## Results

### Expression of inhibitory Siglec receptors is associated with reduced survival in glioma patients

Stratification of glioma patients (combined GBM and low-grade glioma (LGG), The Cancer Genome Atlas, TCGA [22]) by *SIGLEC9* expression (measured with RNA-sequencing) revealed a marked overall survival advantage for patients with lower *SIGLEC9* expression (Fig. 1a). Focusing on all human Siglec receptors in GBM patients alone revealed a significant correlation between high expression and reduced overall survival only for *SIGLEC9* (Supplementary Data Fig. 1a). In contrast, in LGG, expression of 8 out of 15 Siglec receptors correlated significantly with reduced patient survival (Supplementary Data Fig. 1b) [22]. We investigated *SIGLEC9* expression at the single cell level in our single-cell RNA sequencing (scRNA-seq) dataset consisting of five primary GBM patients [23], where we found *SIGLEC9* to be predominantly expressed by GAMs (Fig. 1b). This was also the case for other Siglec receptors showing a negative correlation with overall survival in GBM patients by trend (*SIGLEC1, CD22, SIGLEC7* and *SIGLEC10*) (Supplementary Data Fig. 1a, c). Beside GAMs, we observed expression of *CD22* and *SIGLEC10* in B cells, and of *SIGLEC7* in NK cells, as previously reported by others (Supplementary Data Fig. 1c) [19, 24, 25]. Expression of UDP-GlcNAc 2-epimerase/ManNAc kinase (*GNE*), a rate-limiting enzyme in the sialic acid biosynthesis pathway [26], was mainly expressed by GBM cells (Fig. 1b); suggesting that there could be interactions between Siglec-9 in GAMs and sialic acid in GBM cells. Flow cytometry (FC) analysis of primary human GBM- and glioma-associated mouse MG revealed high expression of Siglec-9 and Siglec-E protein, respectively (Fig. 1c, d, Supplementary Data Fig. 1d, e). A significant upregulation of Siglec-E was observed on mouse MG in the orthotopic tumor context (Fig. 1d). Staining of primary human GBM cells with recombinant Siglec-9 Fc chimeras to determine their sialic acid composition revealed a highly sialylated cell surface (Fig. 1e), which was at similar levels to the mouse malignant astrocytoma cell line CT-2A [27], but not GL261 or the retrovirally induced primary mouse glioma cell line PDGF^+^*Trp53^-^*[28] (Fig. 1f).

**Fig. 1.**
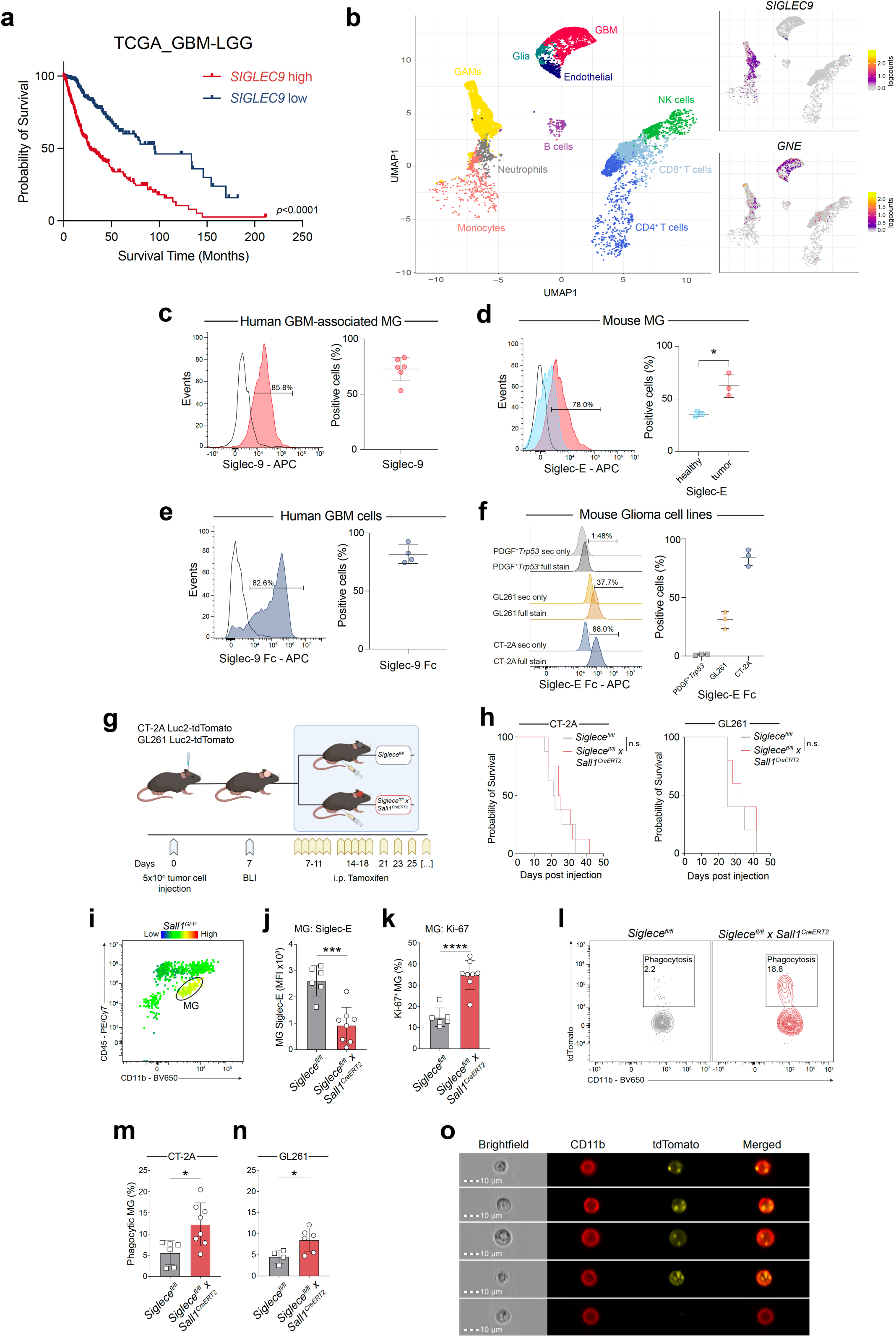
High *SIGLEC9* expression is associated with reduced survival in glioma patients, and its mouse homologue Siglec-E inhibits MG tumor cell phagocytosis. **a,** Kaplan-Meier survival curve of glioma patients (combined GBM and low-grade glioma (LGG)), based on their *SIGLEC9* expression level using the RNA-seq dataset from The Cancer Genome Atlas (TCGA) [22]. The median mRNA expression value was selected as cutoff for high and low expression groups. **b,** Uniform Manifold Approximation and Projection (UMAP) plot showing scRNA-seq cell type annotation in five primary human glioblastoma [23] (left) and UMAP showing expression of *SIGLEC9* and *GNE* in the respective clusters (right). Expression is shown as normalized log_2_ counts. **c, d** Representative histograms and quantification of FC analysis of Siglec-9 expression by human GBM-associated MG (n = 6 donors) **(c)** and Siglec-E expression by mouse MG from healthy and tumor-bearing mice (n = 3 mice per group) **(d) e, f**, Representative histograms and quantification of FC analysis of Siglec-9 ligand expression by human CD45^neg^ GBM cells (n = 4 donors) **(e)** and Siglec-E ligand expression by mouse glioma cell lines derived from PDGF^+^ Trp53^-^ murine gliomas (grey), or cultured GL261 (yellow) and CT-2A (blue) cell lines (n = 3 independent experimental replicates) **(f). g**, Schematic of experimental design **h,** Survival CT-2A (left) and GL261 (right) tumor-bearing animals after MG-conditional Siglec-E deletion (n = 5-8 mice per group). **i,** Gating strategy to identify CD11b^+^CD45^low^ glioma-associated MG in mouse brain tumor single cell suspensions. Gating strategy was confirmed in *Sall1^GFP^ reporter* mice. **j-n,** FC analysis of Siglec-E **(j)** and Ki-67 **(k)** expression in MG. Representative contour plots showing MG CT-2A tumor cell phagocytosis **(l)** and quantification of FC-based MG CT-2A **(m)** and GL261 **(n)** tumor cell phagocytosis measured as percentage of tdTomato^+^ MG (n = 4-8 mice per group). Results shown are from one experiment, representative of two independent experiments **o,** Imaging cytometry showing mouse glioma-associated MG engulfing tdTomato expressing CT-2A tumor cells. Experiment was performed once. Statistics: Data are presented as mean ± SD, log-rank Mantel-Cox test **(a, h)**, unpaired two-tailed Student’s t test **(d, j, k, m, n)**. \**p* ≤ 0.05, \*\*\**p* ≤ 0.001, \*\*\*\**p* ≤ 0.0001.

### Inhibitory Siglec receptors on microglia reduce tumor cell phagocytosis

To investigate the role of Siglec-sialic acid signaling in regulating the MG-mediated anti-tumor immune response, we employed an orthotopic GBM mouse model with MG specific spatio-temporal deletion of *Siglece* by crossing *Siglece^fl/fl^* mice [29] with *Sall1^CreERT2^* mice [30] (*Siglece^fl/fl^* x *Sall1^CreERT2^*). *Sall1^CreERT2^* mice harbor a tamoxifen-inducible *Cre* activity under transcriptional control of the *Sall1* promotor. *Sall1* represents a MG signature gene, not expressed by peripheral MdCs [31]. We induced *Siglece* deletion by intraperitoneal (i.p.) tamoxifen injections beginning 7 days post inoculation of Luc2-tdTomato labelled tumor cells and after confirmation of tumor engraftment by bioluminescence imaging (BLI) (Fig. 1g). We did not observe survival differences in CT-2A or GL261 tumor bearing mice (Fig. 1h), despite efficient deletion of Siglec-E in MG (Fig. 1i, j, Supplementary Data Fig. 1f). Nevertheless, FC analysis of the iTME unveiled high MG proliferation upon Siglec-E knockout (Fig. 1k), accompanied by an enhanced MG GBM-cell uptake measured as the percentage of tdTomato^+^ MG (Fig. 1l-n). The Siglec-E deletion-mediated pro-phagocytic effect in MG was more prominent in CT-2A tumor bearing animals than in GL261-grafted mice (Fig. 1m, n), probably due to the higher Siglec-E ligand expression on CT-2A than in GL261 cells (Fig. 1f). Intracellular uptake of tumor-derived tdTomato fluorescence by MG was microscopically confirmed using imaging flow cytometry (Fig. 1o, Supplementary Data Fig. 1g). Together, these results indicated that inhibitory Siglec-E receptor plays a role in regulating MG-mediated tumor cell phagocytosis. However, perturbing Siglec-E signaling in MG was not sufficient to improve survival, leading us to comprehensively investigate differences between iTME in *Siglece^fl/fl^* and *Siglece^fl/fl^* x *Sall1^CreERT2^*.

### MG activation through Siglec-E deletion induces a counteracting, compensatory MdC response

Co-expression patterns of CD163 and CD86 in infiltrating MdCs (Fig. 2a) revealed a ‘M2-like’ protumorigenic polarization in *Siglece^fl/fl^* x *Sall1^CreERT2^* mice (Fig. 2b, c). Furthermore, we observed a counteracting upregulation of Siglec-E in the total MdC population upon MG-specific Siglec-E deletion (Fig. 2d), particularly prevalent in the more abundant CD163^high^CD86^low^ ‘M2-like’ MdCs (Fig. 2b). Of note, this compensatory upregulation on MdCs was only observed for Siglec-E, whereas we did not measure expression changes in other Siglec receptors (Supplementary Data Fig. 2a).

**Fig. 2.**
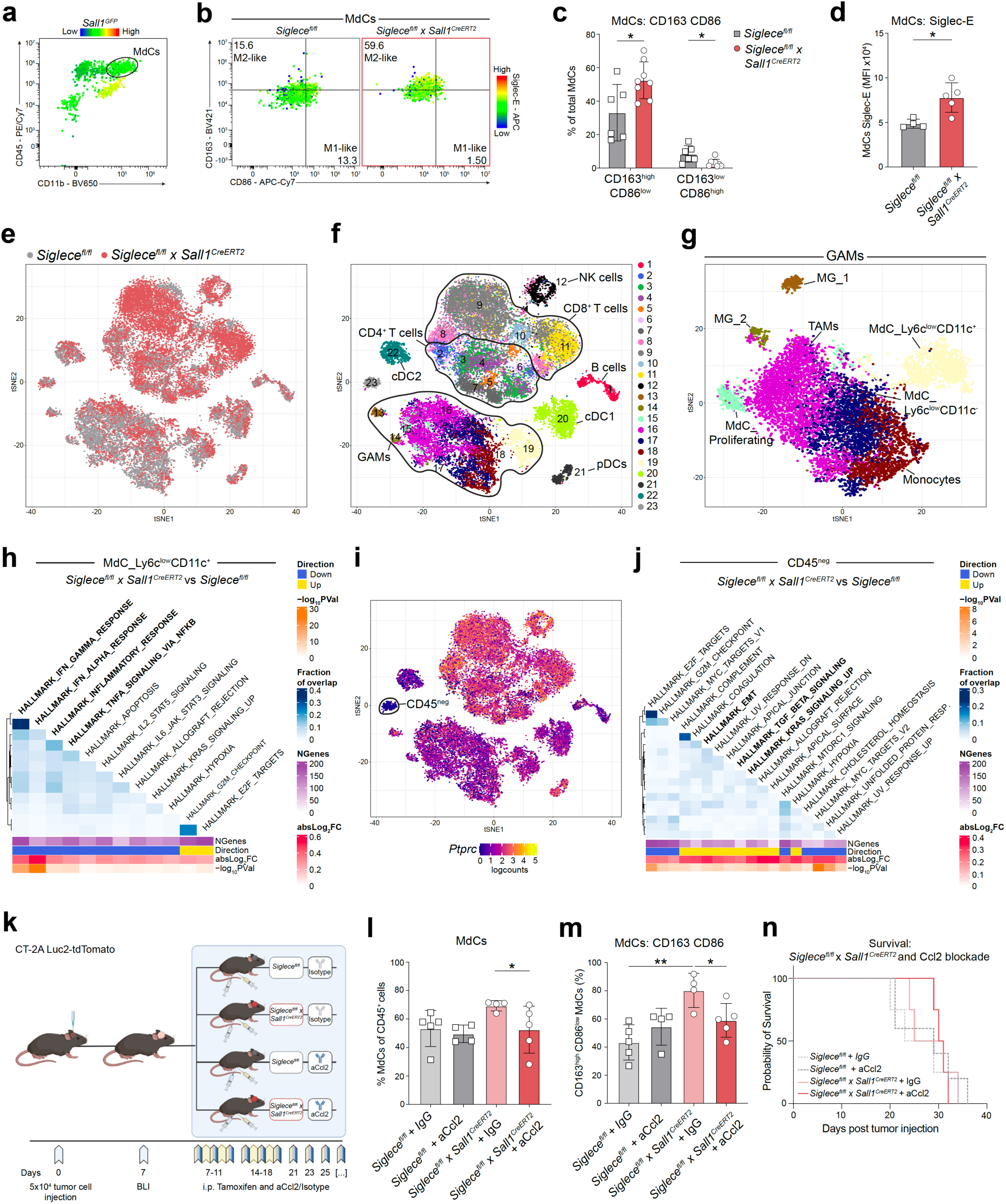
MG activation via MG-specific Siglec-E deletion induces counteracting MdC responses. **a-d**, FC analysis of CD86 and CD163 expression on tumor infiltrating MdCs, identified as CD11b^+^CD45^high^ events. Gating strategy was confirmed in *Sall1^GFP^* reporter mice (**a**), shown as representative dot plots, overlayed with Siglec-E expression (**b**) and quantification of CD163^high^CD86^low^, CD163^low^CD86^high^ co-expression (n = 6-9 mice per group) (**c**) and Siglec-E expression of MdCs (n = 4-5 mice per group) (**d**). Results shown are from one experiment, representative of two independent experiments **e**, t-distributed stochastic neighbor embedding (tSNE) plot from scRNA-seq analysis showing the distribution of sorted immune cells from *Siglece^fl/fl^*(grey) and *Siglece^fl/fl^* x *Sall1^CreERT2^* (red) tumors. **f**, tSNE plot showing the annotated cell populations. **g**, tSNE plot of scRNA-seq analysis showing the subset of GAMs and their annotation. **h**, Heatmap representation of Gene Set Enrichment Analysis (GSEA) results between MdC_Ly6c^low^CD11c^+^ cells from *Siglece^fl/fl^* x *Sall1^CreERT2^* and *Siglece^fl/fl^* tumors using the MSigDB Hallmark collection. The colors on the heatmap represent the fraction of overlap (Jaccard coefficient) between genes annotated to the significant gene sets. NGenes represents the size of the gene sets, and absLog_2_FC represents the average absolute log_2_ fold-change of genes in the gene sets. **i**, tSNE plot showing *Ptprc* (Cd45) expression across cells. Expression is shown as normalized log_2_ counts. **j**, Heatmap representation of GSEA results between CD45^neg^ cells from *Siglece^fl/fl^* x *Sall1^CreERT2^* and *Siglece^fl/fl^* tumors similar to panel (**h**). **k**, Schematic of experimental design. **l**-**n**, FC analysis of tumor infiltrating MdCs from *Siglece^fl/fl^* and *Siglece^fl/fl^* x *Sall1^CreERT2^* CT-2A tumor-bearing mice treated with anti-Ccl2 or isotype control. Percentage of MdCs (**l**), percentage of CD163^high^CD86^low^ co-expressing MdCs (**m**) and survival (**n**) in treatment groups (n = 4-5 mice per group). Experiment was performed once. Statistics: Data are presented as mean ± SD, multiple unpaired Student’s t test with Holm-Sidak’s corrected multiple comparison test (**c**), unpaired two-tailed Student’s t test (**d**), one-way ANOVA with Sidak’s corrected multiple comparison test (**l**, **m**). For statistical analysis of scRNA-seq data, please refer to Methods section. \**p* ≤ 0.05, \*\**p* ≤ 0.01.

Using scRNA-seq, we profiled the iTME from *Siglece^fl/fl^* and *Siglece^fl/fl^* x *Sall1^CreERT2^* CT-2A tumor-bearing mice (Fig. 2e), identifying 23 distinct cell clusters, including seven GAM clusters, ten lymphoid clusters, one NK cell cluster, three dendritic cell (DC) and one B cell cluster (Fig. 2f, Supplementary Data Fig. 2b-d). Focusing on GAM clusters (Fig. 2g), two MG clusters were identified (MG_1 and MG_2). A differential expression analysis between cells from *Siglece^fl/fl^* and *Siglece^fl/fl^* x *Sall1^CreERT2^* CT-2A tumor-bearing mice for these MG clusters revealed a partial reversal of a disease-associated MG (DAM)-phenotype upon Siglec-E deletion (Supplementary Data Fig. 2e). DAMs were initially described in a genetic mouse model of Alzheimer’s disease [32], but have been reported in other models of neurodegeneration and neuroinflammation [33, 34], where excessive activation of MG proinflammatory functions may be detrimental and accelerate the disease [35]. Upon Siglec-E deletion, we observed downregulation of DAM-signature genes [35], including tetraspanins (*Cd9*), chemokines (*Cxcl13*), molecules involved in Trem2-signaling (*Lgals3*) and tissue remodeling (*Spp1, Gpnmb*). In contrast, genes involved in phagocytosis (*Axl*, *Arg1*) and cellular activation (*Trpm2*) were upregulated (Supplementary Data Fig. 2e). Along the same line, gene set enrichment analysis (GSEA) on the differentially expressed genes using the MSigDB Hallmark collection [36] identified TNF-α signaling via NF-κB to be upregulated in Siglec-E depleted MG (Supplementary Data Fig. 2f).

Focusing on MdCs, we identified two phenotypically distinct monocyte-derived cell (MdC) clusters (MdC_Ly6c^low^CD11c^+^ and MdC_Ly6c^low^CD11c^-^), probably representing intermediate stages of differentiation towards monocyte-derived DCs and monocyte-derived macrophages, respectively [37, 38]. Among MdCs clusters, differential expression analysis attributed the highest number of differentially expressed genes (DEGs) to the MdC_Ly6c^low^CD11c^+^ cluster (57 DEGs in MdC_Ly6c^low^CD11c^+^; 12 DEGs in Monocytes, 10 DEGs in MdC_Ly6c^low^CD11c^-^, 12 DEGs in TAMs and 10 DEGs in MdC_Proliferating, at a 5% FDR; Supplementary Table 1). GSEA on the differentially expressed genes ascribed a highly immunosuppressive phenotype to these MdC_Ly6c^low^CD11c^+^ cells upon MG-specific Siglec-E deletion, with downregulation of genes modulating type I and type II interferon responses and TNF-α signaling (Fig. 2h).

In addition, differential abundance analysis revealed significant increases in CD8^+^ T cell clusters 9 and 11 (CD8^+^ T cells_Effector/pre-exhausted and CD8^+^ T cells_Exhausted) in *Siglece^fl/fl^* x *Sall1^CreERT2^* mice (Supplementary Data Fig. 2g). This corroborates the hypothesis that the infiltrating MdCs render the iTME pro-tumorigenic. Although we focused our analysis on immune cells, we still captured transcripts originating from CD45^neg^ cells (Fig. 2i), which acquired a progressive phenotype with upregulation of genes involved in epithelial-mesenchymal transition (EMT), KRAS signaling and increased TGFβ signaling after MG-specific Siglec-E deletion (Fig. 2j), which might facilitate the immunosuppressive shift of the MdC subcluster. Together, these data identified a population of early phase MdCs as a main driver of the counteracting immunosuppressive response upon Siglec-E deletion induced MG activation.

To test our hypothesis of early phase MdCs as main counteracting force to the anti-tumor MG response, we combined MG-specific Siglec-E deletion with C-C chemokine ligand 2 (Ccl2) neutralization (Fig. 2k). This inhibits recruitment of Ccr2-expressing inflammatory monocytes (Supplementary Data Fig. 2c) to the tumor by retaining them in the bone marrow [39]. Indeed, anti-Ccl2 treatment led to less excessive infiltration of MdCs to the tumor site and converted their ‘M2-like’ polarization state upon concomitant MG-specific Siglec-E deletion. However, both phenotypes ameliorated only to control level (Fig. 2l, m), indicating that recruitment of MdCs to the GBM iTME is a highly redundant mechanism which cannot be perturbed by antagonizing the action of a single tumor-attracting chemokine. Accordingly, the combination of MG-specific Siglec E deletion and anti-Ccl2 treatment did not improve survival (Fig. 2n). These results showcase that MG-specific Siglec-E deletion promotes tumor cell phagocytosis, which is subsequently counteracted by infiltrating MdCs that acquire an immunosuppressive phenotype in the perturbed iTME.

### Siglec-E deficiency in whole GAM population improves innate anti-tumor immunity

To test the role of inhibitory Siglec receptors on all GAMs, we used *Cx3cr1^CreERT2^* mice [40], which harbor tamoxifen-inducible *Cre* activity under transcriptional control of the C-X3-C motif chemokine receptor 1 (*Cx3cr1*) promoter. This allowed us to target both GAM populations in the GBM iTME (MG, as well as *Cx3cr1*-expressing MdCs) (Fig. 3a). CT-2A engrafted mice in the *Siglece^fl/fl^* x *Cx3cr1^CreERT2^* group showed a delayed tumor growth measured by in vivo BLI (Fig. 3b, c, Supplementary Data Fig. 3a) resulting in prolonged survival compared to *Siglece^fl/fl^* control mice (Fig. 3d). FC based immune profiling revealed increased MG- and MdC-mediated tumor cell phagocytosis by dual MG/MdC-specific Siglec-E deletion (Fig. 3e, f), accompanied by increased production of effector cytokine TNF-α (Fig. 3f). Additionally, MG displayed accentuated antigen presentation capacity with increased surface MHC-II and costimulatory CD86 expression upon *Cx3cr1*-specific Siglec-E deletion. The MdC compartment showed CD86 increase only by trend, and no difference in MHC-II expression (Fig. 3f). The proportion of intratumoral MdCs was comparable between cohorts at endpoint, indicating that recruitment of these cells to the tumor site remained intact in *Cx3cr1-* specific Siglec-E deleted mice (Supplementary Data Fig. 3b). In fact, MdCs were the dominant CD45^+^ cell population at endpoint (Supplementary Data Fig. 3b), highlighting the potent myeloid influx during CT-2A tumor progression.

**Fig. 3.**
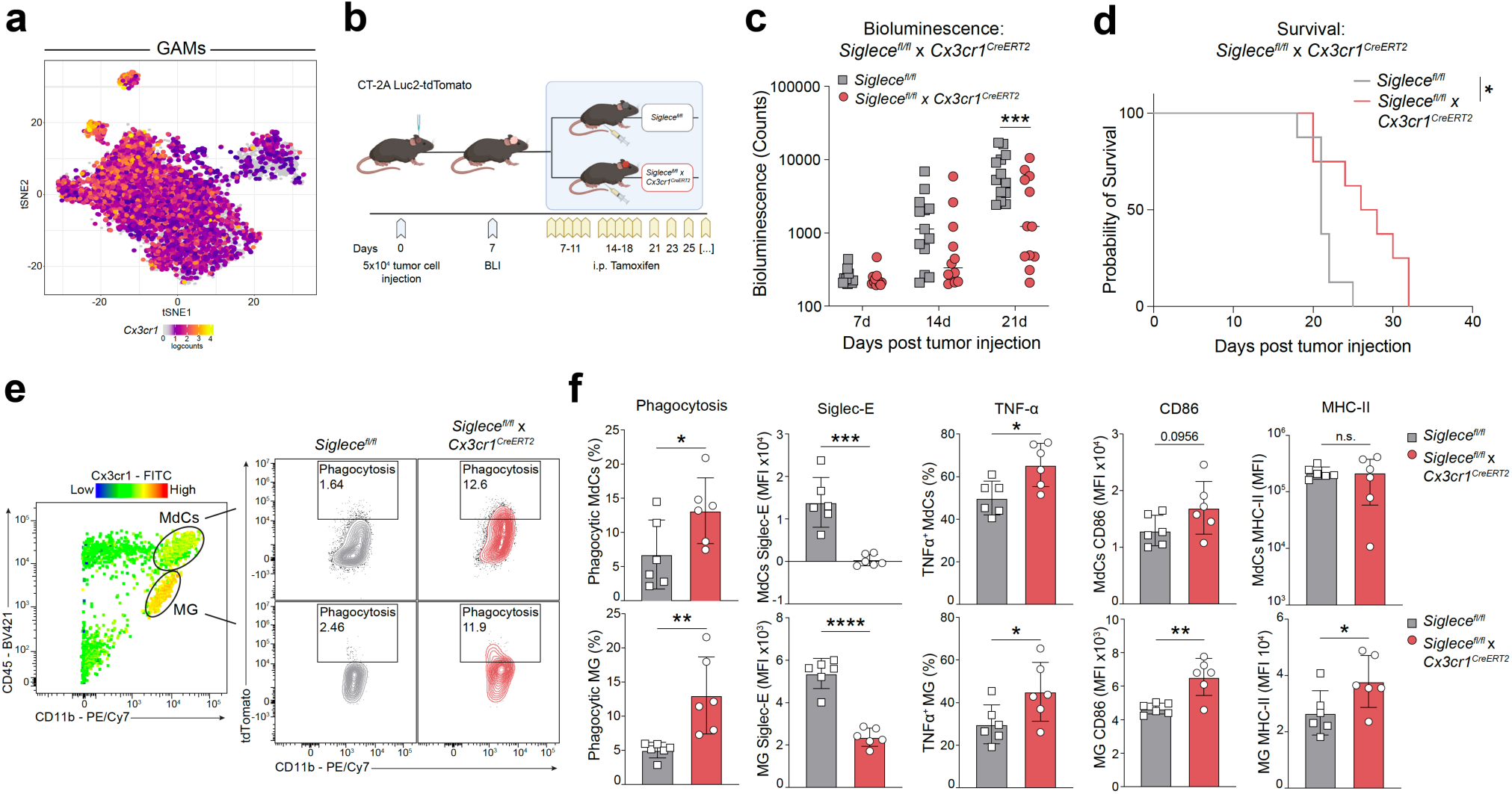
GAM-specific Siglec-E deletion boosts innate anti-tumor immunity. **a**, tSNE plot of scRNA-seq analysis showing *Cx3cr1* expression in the GAM clusters subset. Expression is shown as normalized log_2_ counts. **b**, Schematic of experimental design. **c**, **d**, Surrogate tumor growth assessed by BLI (n = 12-13 mice per group) (**c**) and survival (n = 8 mice per group) (**d**). Results were pooled from two independent experiments. **e**, **f**, FC analysis of MdCs (top row) and MG (bottom row) from *Siglece^fl/fl^* and *Siglece^fl/fl^* x *Cx3cr1^CreERT2^* CT-2A tumor bearing mice. Representative contour plots showing tumor cell phagocytosis assessed by td Tomato^+^ MdCs and MG (**e**) and bar graphs showing quantified phagocytosis as well as expression of Siglec-E, TNF-α, CD86 and MHC-II within MdCs (top row) and MG (bottom row) upon GAM-specific Siglec-E deletion and respective Cre-negative littermate controls (**f**) (n = 6 mice per group). Results shown are from one experiment, representative of two independent experiments. Statistics: Data are presented as median (**c**) and mean ± SD (**f**), two-way ANOVA with Sidak’s corrected multiple comparison test (**c**), log-rank Mantel-Cox test (**d**), unpaired two-tailed Student’s t test (**f**). \**p* ≤ 0.05, \*\**p* ≤ 0.01, \*\*\**p* ≤ 0.001, \*\*\*\**p* ≤ 0.0001.

### Siglec-E deficient MdCs show increased antigen cross-presentation and T cell cross-priming capacity

Within tumor-infiltrating lymphocytes, we observed mainly a CD8^+^ T cell driven response upon GAM-specific Siglec-E deletion with increased activation signature (CD69^+^Ki-67^+^ co-expression) and effector cytokine interferon-γ (IFN-γ) production (Fig. 4a, b). Together with the absent MHC-II response in MdCs after increased tumor cell phagocytosis, we hypothesized that loss of Siglec-E on MdCs could enhance antigen cross-presentation and cross-priming of CD8^+^ T cells. Therefore, we pulsed the CD11b^+^ GAM fraction on day 15 post tumor-inoculation with ovalbumin protein (Fig. 4c) and evaluated the presence of MHC-I bound ovalbumin-derived peptide SIINFEKL. Indeed, we noted a significant increase in antigen cross-presentation by MdCs, but not MG, upon Siglec-E deletion (Fig. 4d). Co-culture of ovalbumin-pulsed CD11b^+^ GAMs with naive CD8^+^ T cells from OT-I transgenic mice expressing ovalbumin-specific T cell receptors, resulted in increased OT-1 T cell activation in an antigen-specific manner by GAMs from *Siglece^fl/fl^* x *Cx3cr1^CreERT2^* mice compared to *Siglece^fl/fl^* control (Fig. 4e-g). In vivo depletion of CD8^+^ T cells using an anti-CD8 antibody diminished the anti-tumor effect of GAM Siglec-E deletion (Fig. 4h, i). Of note, we observed a delayed tumor growth in the CD8^+^ T cell depleted *Siglece^fl/fl^* x *Cx3cr1^CreERT2^* mice compared to *Siglece^fl/fl^* until day 14 post tumor injection (Fig. 4h), confirming increased level of innate immune tumor control of Siglec-E deletion during early tumor growth. However, Siglec-E deletion-driven tumor cell phagocytosis seems insufficient to induce a lasting anti-tumor immune response which ultimately relies on the generation of adaptive immunity mediated by cytotoxic CD8^+^ T cells.

**Fig. 4.**
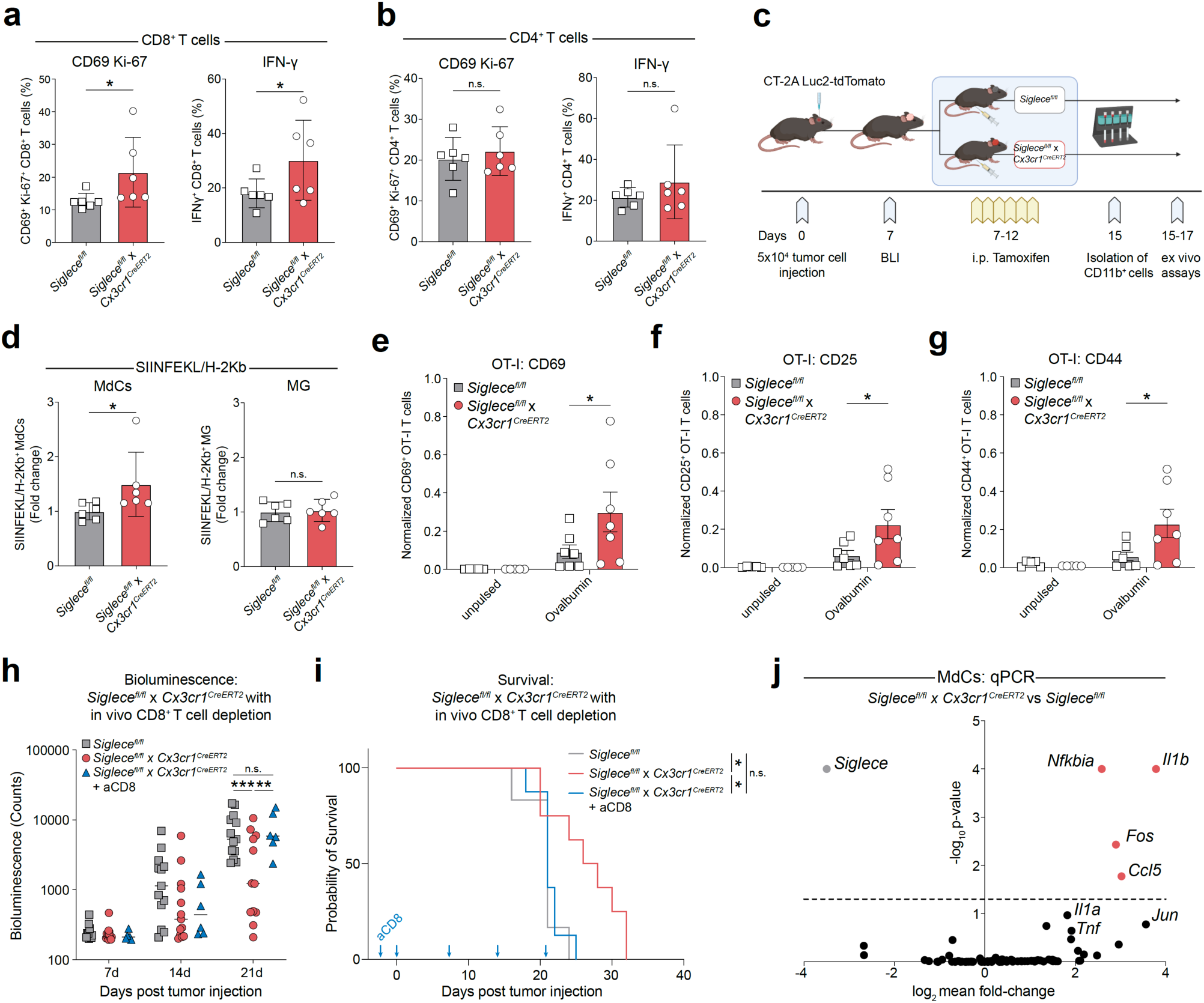
Intratumoral Siglec-E-deleted MdCs show increased antigen cross-presentation and T cell cross-priming capacity ex vivo. **a**, **b**, FC analysis of CD69 and Ki-67 co-expression (left panels) and intracellular IFN-γ production (right panels) in tumor-infiltrating CD8^+^ (**a**) and CD4^+^ T cells (**b**) from *Siglece^fl/fl^* and *Siglece^fl/fl^* x *Cx3cr1^CreERT2^* CT-2A tumor-bearing animals (n = 6 mice per group). Results shown are from one experiment, representative of two independent experiments. **c**, Schematic of experimental design. **d**, FC analysis of SIINFEKL peptide bound to H-2Kb on MdCs (left) and MG (right) (n = 6 mice per group). Results were pooled from two independent experiments. **e**-**g**, FC analysis of CD69 (**e**), CD25 (**f**) and CD44 (**g**) on SIINFEKL peptide-specific OT-I T cells, after co-culture with unpulsed or ovalbumin-pulsed CD11b^+^ GAMs (n = 7 mice per group). Results were pooled from two independent experiments. **h**, **i**, BLI as surrogate for tumor growth (n = 6-13 mice per group) (**h**) and survival (n = 6-8 mice per group) (**i**) of in vivo CD8^+^ T cell depleted *Siglece^fl/fl^* x *Cx3cr1^CreERT2^* CT-2A tumor-bearing animals, compared to animals from Fig. 3c and 3d. Blue arrows in Fig. 4i indicate anti-mouse CD8a i.p. administrations (days −2, 0, 7, 14 and 21). Results shown are from one experiment, representative of two independent experiments. **j**, qPCR analysis of NF-κB target genes in MdCs sorted from *Siglece^fl/fl^* and *Siglece^fl/fl^* x *Cx3cr1^CreERT2^* CT-2A tumor-bearing mice on day 15 after tumor cell injection, and 8 days after induction of GAM-specific Siglec-E deletion (n = 9-10 mice per group). Results were pooled from three independent experiments, with n = 3-4 mice pooled per genotype each. qPCR analysis of *Siglece* was done separately, n = 4 mice per genotype. Statistics: Data are presented as mean ± SD (**a, b, d**), mean ± SEM (**e-g**) and median (**h**), unpaired Student’s t test (**a**, **b**), two-tailed Mann-Whitney test (**d**), one-way ANOVA with Sidak’s corrected multiple comparison test (**e**-**g**), two-way ANOVA with Sidak’s corrected multiple comparison test (**h**), log-rank Mantel-Cox test (**i**), two-way ANOVA for NF-κB target genes and unpaired two-tailed Student’s t test for *Siglece* (**j**). \**p* ≤ 0.05, \*\**p* ≤ 0.01, \*\*\**p* ≤ 0.001.

Efficient priming of CD8^+^ T cells and promoting their effector functions requires both antigen cross-presentation and secretion of pro-inflammatory cytokines by MdCs [41]. An unbiased proteomic analysis of tumor-associated MdCs revealed overrepresentation of the IL-17 signaling pathway in Siglec-E deleted compared to wildtype MdCs, using the KEGG Pathway database (Supplementary Data Fig. 4b) [42]. IL-17 is known to trigger activation of the canonical nuclear factor κB (NF-κB) cascade, and subsequently upregulates expression of various pro-inflammatory genes [43]. To test this, we profiled signature genes within the NF-κB signaling pathway in tumor-associated MdCs 15 days post tumor engraftment by quantitative real-time PCR (qPCR). This revealed increased expression of pro-inflammatory genes (*Il1b, Ccl5* and *Tnf, Il1a* by trend) and genes associated with the activator protein 1 transcription-complex (AP-1) (*Fos* and *Jun* by trend) in MdCs from *Siglece^fl/fl^* x *Cx3cr1^CreERT2^* mice versus *Siglece^fl/fl^* animals (Fig. 4j, Supplementary Data Fig. 4c). Increased NF-κB signaling induced also negative feedback circuits by increased expression of *Nfkbia* which is one of the earliest genes induced following NF-κB activation leading to termination of NF-κB signaling [44]. Collectively, these findings indicate that GAM-specific Siglec-E deletion promotes tumor cell phagocytosis by MG and MdCs, enhances intratumoral CD8^+^ T cell responses by antigen cross-presentation and increases production of pro-inflammatory cytokines by MdCs, potentially mediated by NF-κB signaling axis.

### Genetic targeting of sialic acid on CT-2A cells recapitulates main findings of GAM-specific Siglec-E deletion in vivo

To complement the effect of immune evasion mediated by Siglec-E receptor, we targeted its sialic acid ligands on CT-2A cells by knocking out the GNE enzyme (CT-2A^ΔGNE^) (Fig 5a, Supplementary Data Fig. 5a). CT-2A^ΔGNE^ cells showed no differences with regard to their in vitro proliferation and viability compared to wildtype control (CT-2A^WT^) (Supplementary Data Fig. 5b). Orthotopic transplantation of CT-2A^ΔGNE^ cells into C57BL/6 mice resulted in increased tumor cell phagocytosis, TNF-α production and a MG-restricted MHC-II response (Fig. 5c), recapitulating the main findings from the GAM Siglec-E deletion in both innate immune populations. Although we observed a favorable CD4^+^ T cell response with less CD25^+^Foxp3^+^ Tregs and more Cxcr3^+^T-bet^+^ Th1 CD4^+^ T cells in CT-2A^ΔGNE^ tumor bearing animals at endpoint (Supplementary Data Fig. 5c), the main driver of the adaptive immune response after innate immune activation were CD8^+^ T cells. This was exemplified by less abundant PD-1^+^, TIM-3^+^, LAG-3^+^ and CTLA-4^+^ co-expressing CD8^+^ T cells, greater degranulation capacity and increased IFN-γ production in CT-2A^ΔGNE^ tumor bearing animals (Fig. 5d, e), ultimately improving survival (Supplementary Data Fig. 5d). Together, targeting Siglec receptor ligands in the tumor recapitulates the main findings of GAM-specific Siglec-E deletion in the host.

**Fig. 5.**
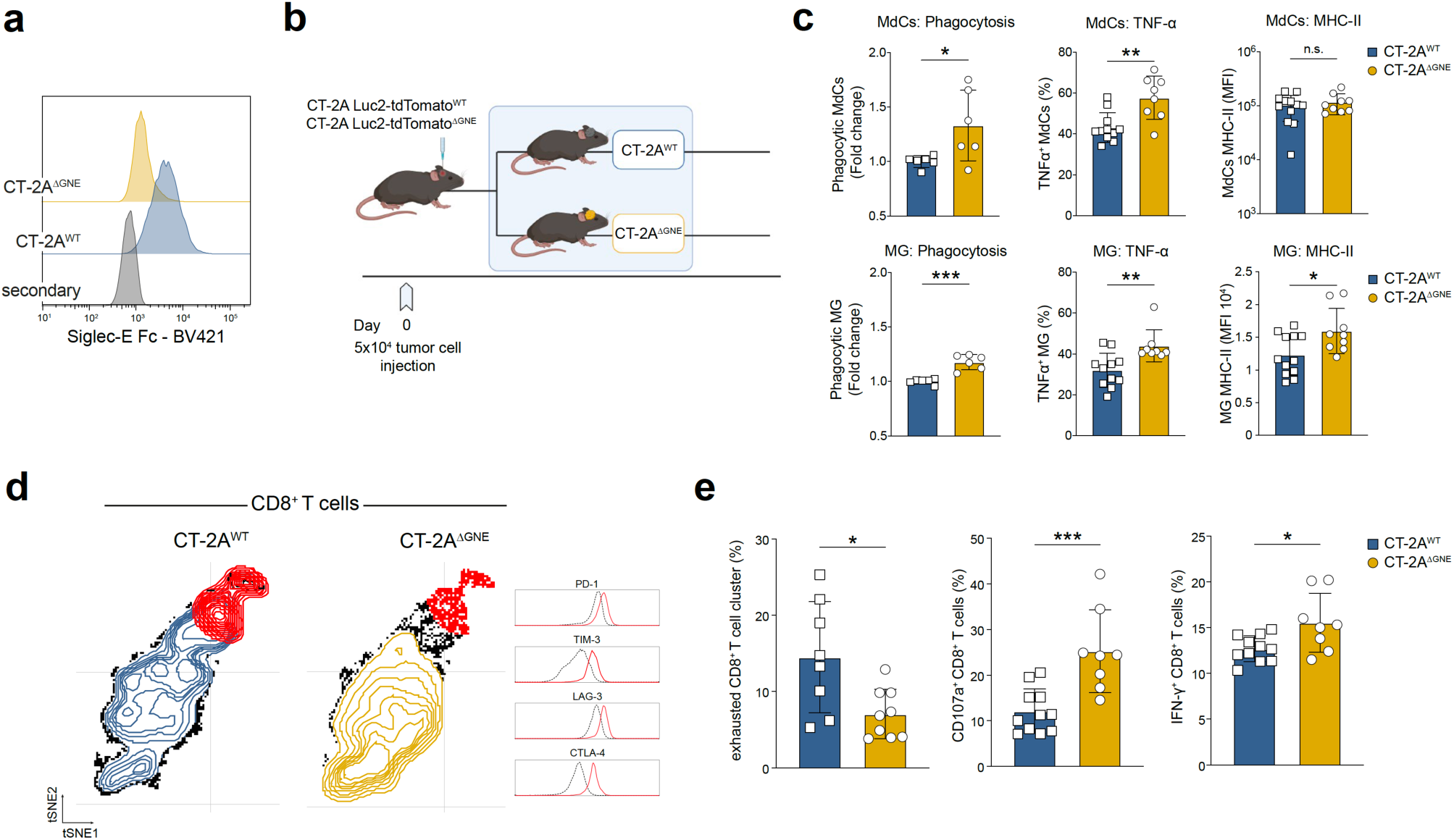
Genetic targeting of sialic acid on CT-2A cells recapitulates main findings of Siglec-E deletion in the host. **a**, Representative histogram of Siglec-E Fc staining on CT-2A^WT^ and CT-2A^ΔGNE^ cells. **b,** Schematic of experimental design. **c**, FC analysis of phagocytosis, and TNF-α and MHC-II expression on MdCs (top row) and MG (bottom row) from CT-2A^WT^ and CT-2A^ΔGNE^ injected C57BL/6 wildtype mice (n = 6 mice per group for phagocytosis, n = 8-11 mice per group for other analyses). Results were pooled from two independent experiments. **d**, FC analysis of inhibitory T cell receptors on intratumoral CD8^+^ T cells visualized on tSNE map (tSNE maps show concatenated CD8^+^ T cells from n = 8 mice per group). Red cluster marks exhausted CD8^+^ T cells identified by high co-expression of inhibitory receptors (PD-1, TIM-3, LAG-3 and CTLA-4), indicated by histograms. Red histogram marks median marker expression in red cluster (exhausted CD8^+^ T cell cluster) and black dotted histogram indicates median marker expression in the remaining cells. **e**, Quantification of FC analysis, showing percentage of exhausted CD8^+^ T cells (PD-1^high^, TIM-3^high^, LAG-3^high^ and CTLA-4^high^) (left), CD107a^+^ (middle) and IFN-γ^+^ (right) intratumoral CD8^+^ T cells between CT-2A^WT^ and CT-2A^ΔGNE^ injected C57BL/6 wildtype mice (n = 8-11 mice per group). Results were pooled from two independent experiments. Statistics: Data are presented as mean ± SD, unpaired two-tailed Student’s t test. \**p* ≤ 0.05, \*\**p* ≤ 0.01, \*\*\**p* ≤ 0.001.

### GAM-specific Siglec-E deletion improves combinatorial immunotherapy in GBM

To harness and further elucidate the therapeutic potential of GAM-specific Siglec-E deletion (*Siglece^fl/fl^* x *Cx3cr1^CreERT2^*), we initiated combination treatments with CD47 blockade (Fig.6a), an established innate immunotherapeutic agent [8–10]. It has been previously shown that CD47 blockade enhances tumor cell phagocytosis and T cell cross-priming as well [45]. We observed a significant reduction of tumor growth and improved overall survival in the combinatorial condition with 11% (1/9) of animals showing tumor rejection (Fig. 6b, c). In *Siglece^fl/fl^* x *Cx3cr1^CreERT2^* tumors collected at endpoint, tumor-infiltrating CD8^+^ T cells demonstrated high level of PD-1 expression, while PD-L1 was significantly upregulated on CD45^neg^ and CD45^pos^ cells (Fig. 6d, e). This compensatory T cell checkpoint upregulation could be caused by increased CD8^+^ T cell activation and IFN-γ production after GAM-specific Siglec-E deletion (Fig. 4a) [46]. To overcome this potential resistance mechanism, we additionally treated tumor bearing *Siglece^fl/fl^* x *Cx3cr1^CreERT2^* mice with both CD47 and PD-1 blocking antibodies (Fig. 6f). These triple-therapy treated animals exhibited a sustained survival benefit, with 23% (3/13) of animals having tumor rejection (Fig. 6g, h). Contra-lateral hemisphere tumor-rechallenging of surviving animals in this treatment cohort led to tumor rejection, whereas GAM-Siglec-E deleted/anti-CD47 treated mice succumbed to tumor progression after rechallenge (Fig. 6i). This data suggests that development of a lasting immunological memory and tumor control in GBM can only be achieved by targeting both innate and adaptive immune checkpoints to tackle the manifold immune tolerance mechanisms occurring during tumor progression.

**Fig. 6.**
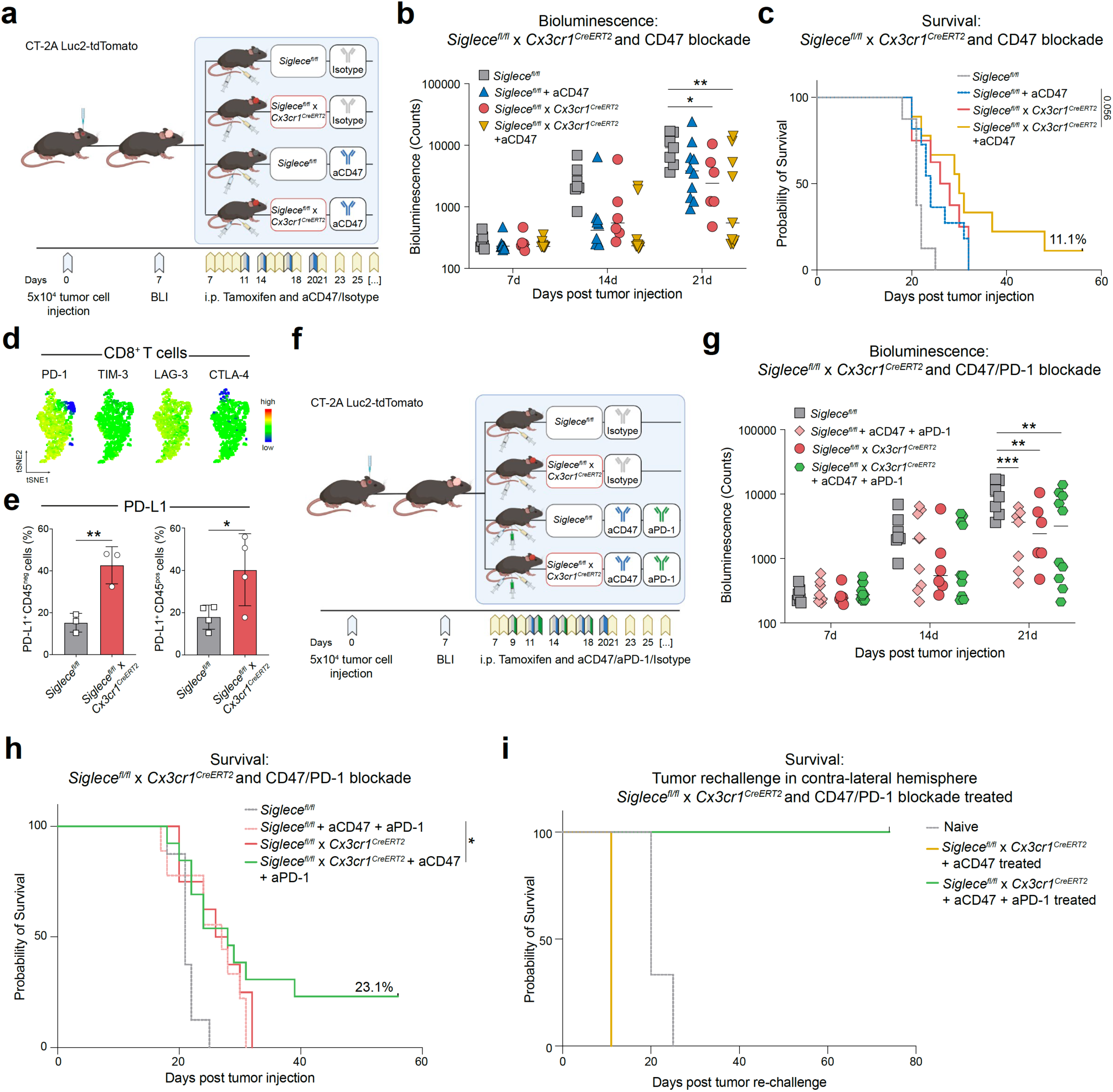
Siglec-E deletion boosts GBM immunotherapy. **a**, Schematic of experimental design. **b**, **c**, Surrogate tumor growth assessed by BLI (n = 6-10 mice per group) (**b**) and survival (n = 8-11 mice per group) (**c**) of *Siglece^fl/fl^* and *Siglece^fl/fl^* x *Cx3cr1^CreERT2^* CT-2A tumor-bearing animals treated with anti-CD47 or isotype control. Results were pooled from two independent experiments. **d**, FC analysis of inhibitory T cell receptors on intratumoral CD8^+^ T cells shown on tSNE map (tSNE maps shows concatenated CD8^+^ T cells from n = 6 mice per group) **e**, FC analysis of PD-L1 expression on CD45^neg^ (left) and CD45^pos^ cells (right) from of *Siglece^fl/fl^* and *Siglece^fl/fl^* x *Cx3cr1^CreERT2^* mice (n = 3-4 mice per group). **f**, Schematic of experimental design. **g**, **h**, Surrogate tumor growth assessed by BLI (n = 6-10 mice per group) (**g**) and survival (n = 8-13 mice per group) (**h**) of *Siglece^fl/fl^* and *Siglece^fl/fl^* x *Cx3cr1^CreERT2^* CT-2A tumor-bearing animals treated with anti-CD47 and anti-PD1 or isotype control. Results were pooled from two independent experiments. **i**, Rechallenge of the tumor-free mice from Fig. 5c and 5h and tumor-naive control mice with intracranial injection of 5 x 10^4^ CT-2A tumor cells into the contra-lateral hemisphere (n = 1 for *Siglece^fl/fl^* x *Cx3cr1^CreERT2^* + aCD47 treated, n = 3 mice per group for other two groups). Statistics: Data are presented as median (**b, g**) and mean ± SD (**e**), two-way ANOVA with Sidak’s corrected multiple comparison test (**b**, **g**), unpaired two-tailed Student’s t test (**e**), restricted mean survival time (RMST) comparison (**c, h**). \**p* ≤ 0.05, \*\**p* ≤ 0.01, \*\*\**p* ≤ 0.001, \*\*\*\**p* ≤ 0.0001.

### Siglec-9 blockade induces immune response and anti-tumor activity in human GBM explants

To determine the translational potential of Siglec-9 blockade, we prospectively collected GBM specimens from four primary and one recurrent GBM patient undergoing neurosurgical resection. All samples were neuropathologically diagnosed as GBM grade 4, *Isocitrate Dehydrogenase 1/2* (*IDH*) wild type (Supplementary Table 2). Intact tumor fragments (explants) were subsequently cultured in 3D perfusion bioreactors for five days in the presence or absence of Siglec-9 blocking antibody (Fig. 7a). As previously reported by us, this culture system provides flow of the media through the tissue, enabling culturing intact tissue of greater thickness and thereby better preserving the GBM iTME compared to static conditions (Fig. 7b) [47].

**Fig. 7.**
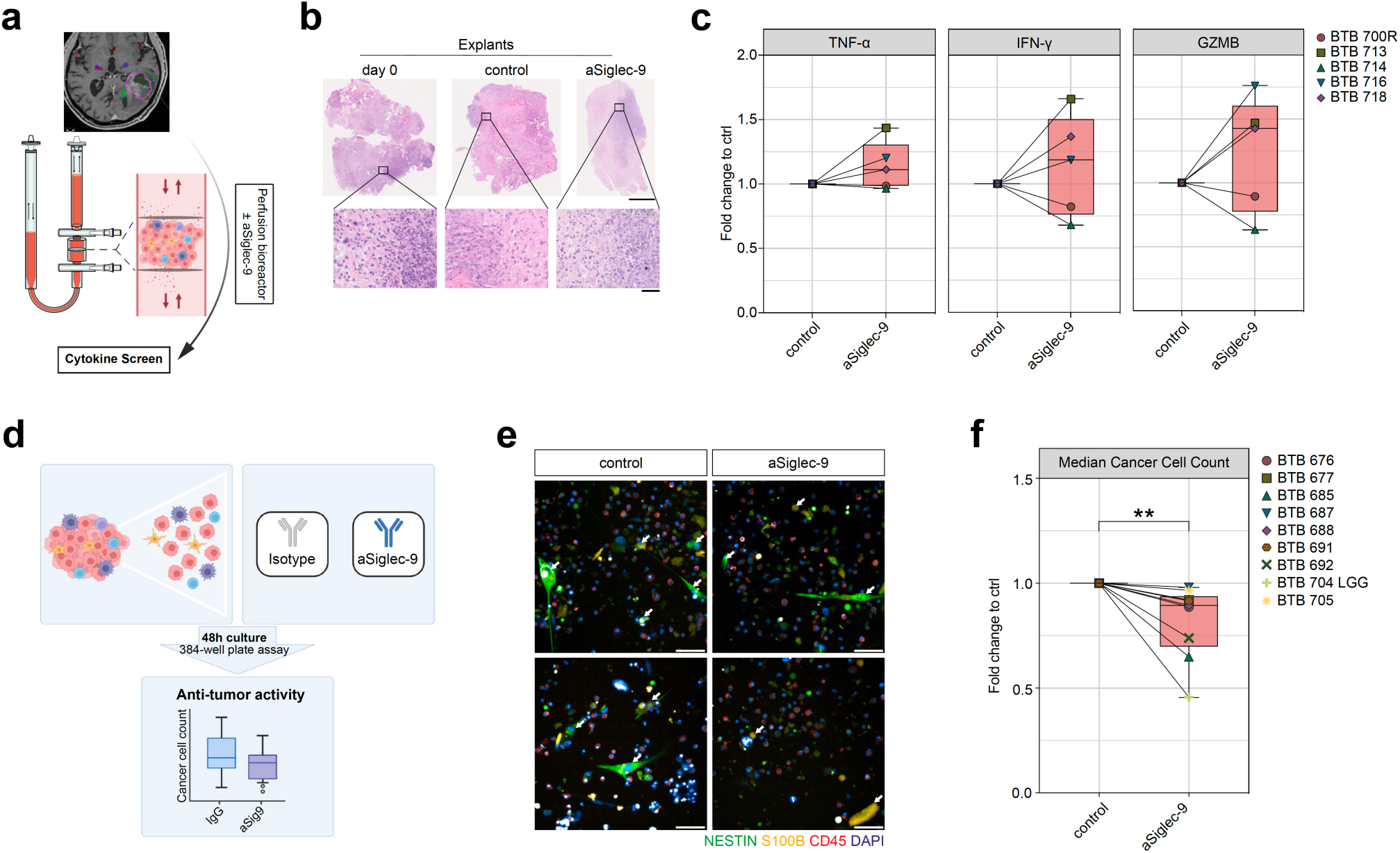
Siglec-9 blockade induces immune response and anti-tumor activity in human GBM explants. **a**, Schematic of experimental design. Fresh tumor biopsies were taken and directly transferred into 3D perfusion bioreactors. Explants were cultured for 5 days in the presence or absence of anti-Siglec-9 blocking antibody. Soluble proteins from bioreactor media were measured by PEA to assess response per patient sample. **b**, Representative H&E-stained images of formalin-fixed paraffin-embedded explants on the day of tumor resection (day 0) and after 5 days of culture in perfusion bioreactors. Scale bars, 1000 μm (overview) and 50 μm (close-up). **c**, Fold change in TNF-α, IFN-γ and GZMB secretion measured in the media of anti-Siglec-9 treated versus control bioreactors, for each individual patient. **d**, Schematic of experimental design. Glioma samples (n = 9) were dissociated and cultured for 48 hours in the presence of Siglec-9 blocking antibody or isotype control. **e**, Representative immunofluorescence images of one patient sample (BTB 688) treated with isotype (left) or anti-Siglec-9 (right). White arrows denote cancer cells. Scale bar, 50 μm**. f**, Fold change in median cancer cell count in anti-Siglec-9 treated versus control. Per patient, 6 technical replicates for control and 3 technical replicates for anti-Siglec-9 samples (Supplementary Data Fig 6b). Median of technical replicates is used and plotted as fold change. Statistics: unpaired two-tailed Student’s t test (**f**). **p ≤ 0.01.

Analysis of secreted soluble proteins by highly sensitive proximity extension assay (PEA) technology after 5 d in culture identified three out of five patients (60%) as responders to Siglec-9 blockade, as indicated by a signature of induced TNF-α, IFN-γ and Granzyme B (GZMB) expression (Fig. 7c). Notably, one of the non-responders was the patient with recurrent GBM (BTB 700R). Within the responders, the observed increase was significant for IFN-γ and Granzyme B and by trend for TNF-α (*p* = 0.06) (Supplementary Data Fig. 6a).

Next, we assessed the anti-tumor activity of Siglec-9 blockade-induced immune activation on a single cell level. Single cell suspensions from nine additional newly diagnosed glioma patients (eight GBM, grade 4, *IDH* wildtype and one LGG; Supplementary Table 2), were exposed to Siglec-9 blocking antibody or control for 48 h (Fig. 7d). Using an automated image-based screening platform [48] to read-out S100B^+^CD45^-^ or NESTIN^+^CD45^-^ glioma cell counts (Fig. 7e), Siglec-9 disruption efficiently reduced the number of glioma cells despite interpatient heterogeneity (Fig. 7f, Supplementary Data Fig. 6b).

## Discussion

Here, we identified the Siglec-sialic acid-axis as innate immune inhibitory pathway in GBM mediating a strongly immunosuppressive TME. We demonstrate that the deletion of Siglec receptors on MG and MdCs or reduction of Siglec ligands on tumor cells can reverse this immune suppression allowing successful combinatorial immunotherapy in preclinical models. As the main mechanism, we show that Siglec-E deletion leads to increased tumor cell phagocytosis by MG and MdC and elevated expression of NF-κB target genes in MdCs. This mediates cross-priming of CD8^+^ T cells and ultimately synergizes with cancer immunotherapy to convey a substantial survival benefit in a highly aggressive and poorly immunogenic CT-2A GBM preclinical model [49].

Previous preclinical studies have shown similar anti-phagocytic [17, 50–52] and macrophage differentiating properties [15, 16, 53] for Siglec receptors in cancer and other diseases. Our study expands this knowledge on the interactions of Siglec receptors with sialoglycans by dissecting the complex interplay between the two main innate immune populations in the GBM iTME. We found a counteracting MdC response upon Siglec-E deletion-driven MG activation and proliferation. Yeo and colleagues recently reported similar changes in their study investigating longitudinal changes in the immune cell composition throughout tumor progression in a genetic mouse GBM model. [54] Specifically, they identified a highly proliferating population of GBM-associated MG, for which the authors discussed a decisive role in activating emergency myelopoiesis in GBM and recruiting bone marrow-derived immunosuppressive myeloid cells to the GBM iTME [54]. This paralleled our observation of a Siglec-E deletion induced activation and proliferation of MG cells, and the counteracting ingress of immunosuppressive MdCs which could only be reverted to control level by Ccl2 inhibition. Even though, the tumor cells were not the primary focus of our scRNA-seq analysis after MG Siglec-E disruption, we found profound changes in the CT-2A transcriptome as well, particularly concerning EMT pathways. This might unveil further tumor-cell intrinsic plasticity and resistance mechanisms upon perturbance of iTME components e.g., MG, and highlights potential paracrine and intercellular reactions between neoplastic cells and MG induced by highly selective deletion of Siglec-E.

By extending the cell type specific Siglec-E deletion to the MdC compartment, we observed increased anti-tumor immunity and upregulation of NF-κB target genes. Others similarly attributed a role as a negative modulator of NF-κB activity to Siglec-9 [51, 55]. In our RNA validation data, *Ccl5* was among the highest upregulated NF-κB target genes upon MdC-specific Siglec-E deletion. Several studies showed positive correlations between expression of inflammatory chemokine Ccl5 and immune cell recruitment to the tumor [56–58]. However, some controversy arose regarding the role of Ccl5 in cancer, as other studies suggested that Ccl5 has potential tumor-promoting effects, either by directly affecting tumor growth by expanding cancer stem cells [59] or promoting immune escape by stabilizing PD-L1 [60]. Unlike IFN-γ, which enhances PD-L1 expression at the transcriptional level [46], Ccl5 has been shown to modulate deubiquitination and stability of PD-L1 [60]. This might contribute to the adaptive resistance after Siglec-E deletion, and illustrate the complex relationship between innate and adaptive immune responses to build an effective and persistent antitumor immunity.

Our data highlight the Siglec-sialic acid axis as an attractive therapeutic target in GBM patients. Together with recent findings, our study further underlines the importance of combining innate and adaptive immunotherapies, especially in less immunogenic and ICI-resistant tumors such as GBM [45]. Targeting sialic acids as ligands for Siglec receptors on tumor cells represents an alternative approach to therapeutically disrupt the Siglec-sialic acid pathway, as demonstrated by genetically targeting sialic acid biosynthesis in CT-2A cells. By applying this strategy, concerns regarding functional redundancy and potential compensatory mechanisms after blockade of one Siglec receptor would be mitigated. In line with this, recent work showcased high efficacy of tumor cell desialylation [17, 52, 61–63]. However, even a targeted approach for example by using antibody-sialidase conjugates would most likely cause severe adverse events, given that sialic acid participates as an integral part of ganglioside structure in synaptogenesis and neural transmission [64]. Additional work will be needed to identify GBM cell-specific sialylation patterns enabling cancer cell targeted desialylation therapies.

Using cultured, perfused 3D tumor explants (as recently described by us [47]) and an image-based ex-vivo drug screening platform, we showcased the translational relevance of Siglec-9 disruption within GBM. Notably, we observed the biggest decrease in tumor cell count after Siglec-9 blockade in the LGG patient and considering the TCGA survival data, further studies on the role of Siglec receptors in LGG are needed.

Taken together, we show that loss of inhibitory Siglec receptors promotes glioma-associated MG and MdCs to phagocytize GBM cells and improve cross-presentation and subsequent T cell activation. Using a poorly immunogenic GBM preclinical model, we demonstrated the synergistic therapeutic potential of combined Siglec-E blockade with ICI against GBM to facilitate innate and adaptive anti-tumor immune responses. Furthermore, we demonstrated the translational potential of Siglec-9 blockade-induced immune activation in patient-derived explant cultures, paving the way to local therapy regimens. These results build on growing interest in designing combination immunotherapies with innate and adaptive ICI, and underscore the value of Siglec blockade in liberating innate immune responses to potentiate anti-tumor immunity.

## Methods

### Human samples

Human adult GBM tissue samples were obtained from the Neurosurgical Clinic of the University Hospital of Basel, Switzerland in accordance with the Swiss Human Research Act and institutional ethics commission (EKNZ 02019-02358). All patients gave written informed consent for tumor biopsy collection and signed a declaration permitting the use of their biopsy specimens in scientific research, including storage in our brain tumor biobank (Req-2019-00553). All patient identifying information was removed and tissue was coded for identification. Patient characteristics from all participating patients are listed in Supplementary Table 2.

### Human GBM tissue dissociation

Resected GBM tissue samples were immediately placed on ice and transferred to the laboratory for single cell dissociation within 2-3 h after resection. Human brain tissue was manually minced using razor blades and enzymatically dissociated at 37°C for 30 minutes with 1 mg mL^-1^ collagenase-4 (#LS004188, Worthington Biochemical Corporation, USA) and 250 U mL^-1^ DNAse1 (#10104159001, Roche, Switzerland) in a buffer containing Hank’s Balanced Salt Solution (HBSS) with Ca^2+^/Mg^2+^, 1% MEM non-essential amino acids (Gibco), 1 mM sodium pyruvate (Gibco), 44 mM sodium bi-carbonate (Gibco), 25 mM HEPES (Gibco), 1% GlutaMAX (Gibco) and 1% antibiotic-antimycotic (Sigma-Aldrich). Cells were filtered and separated from dead cells, debris and myelin by a 0.9 M sucrose (#84100, Simga Aldrich, USA) density gradient centrifugation. Upon ACK-lysis for removal of erythrocytes (Gibco) the single-cell suspension was used for flow cytometry.

### Mice

All animal handling, surveillance, and experimentation were performed according to the guidelines and legislation of the Swiss Federal Veterinary Office (SFVO) and the Cantonal Veterinary Office, Basel-Stadt, Switzerland, under license # 2929. *Siglece^fl/fl^* [29] mice were crossed to *Sall1^CreERT2^* [30], kindly provided by Prof. Nishinakamura (Kumamoto University, Kumamoto, Japan) or to *Cx3cr1^CreERT2^* [40] mice, received from Prof. Niess (University Hospital Basel, Basel, Switzerland) generating offspring mice harboring a tamoxifen-inducible Siglec-E conditional knockout in MG or GAM population, respectively. For both *Cre* lines *Siglece^fl/fl^* mice were used as controls. Rag2/OT-I (*Rag2^tm1Fwa^* Tg(TcraTcrb)1100Mjb) mice were kindly provided by Prof. Alfred Zippelius (University Hospital Basel, Basel, Switzerland) and originally obtained from Taconic (USA). C57BL/6 mice were obtained from Janvier Labs (France) and bred in-house at the Department of Biomedicine, University Hospital Basel, Basel, Switzerland. Mice of both sexes were used throughout the study; sex-matched and age-matched controls were used in individual experiments. All animals were euthanized before reaching humane endpoint, including loss of locomotor activity, weight loss (up to 20%) and central nervous system symptoms. The survival time was measured from the day of tumor cell implantation to the day of euthanasia. Mice of different treatment groups were co-housed in the same cage to blind experimenter in determining humane endpoint.

### Mouse Genotyping

The following primers were used: *Sall1^CreERT2^* (1): AGC TAA AGC TGC CAG AGT GC; *Sall1^CreERT2^* (2): CAA CTT GCG ATT GCC ATA AA; *Sall1^CreERT2^* (3): GCG TTG GCT ACC CGT GAT AT. *Cx3cr1^CreERT2^* (1): AAG ACT CAG GTG GAC CTG CT; *Cx3cr1^CreERT2^* (2): CGG TTA TTC AAC TTG CAC CA; *Cx3cr1^CreERT2^* (3): AGG ATG TTG ACT TCC GAG TTG. *Siglece^fl/fl^* (1): CAG CCC ATC TTT GGC AGA TCC TTG T; *Siglece^fl/fl^* (2): AGT CAA AAC AAA CAC AGC ACA AGC C. PCR amplification was done using GoTaq G2 Green Master Mix (#M7823, Promega, USA).

### Cell lines

The mouse malignant glioma cell line GL-261 was purchased from the DSMZ-German Collection of Microorganisms and Cell Cultures GmbH, Leibniz-Institute (Braunschweig, Germany). The mouse malignant astrocytoma cell line CT-2A was a kind gift from Prof. Seyfried (Boston College, Chestnut Hill, USA). The murine cell lines were cultured in Dulbecco’s modified Eagle medium (DMEM), high glucose, no glutamine (Gibco) supplemented with 10% fetal bovine serum (FBS) (PAN-Biotech), 1% GlutaMAX (Gibco) and 1% Penicillin-Streptomycin (Sigma-Aldrich). All tumor cell lines were cultured as adherent monolayers, maintained at 37°C, 5% CO_2_ and regularly split when reached around 80% confluence. All cell lines were regularly tested for mycoplasma contamination using a biochemical test kit (#LT07-318, Lonza, Switzerland) and were free of mycoplasma contamination.

### Transduction of mouse GBM cell lines

For in vivo tumor monitoring by bioluminescence imaging (BLI) and fluorescent labelling of tumor cells in FC-based phagocytosis assay, GL261 and CT-2A cells were lentivirally transduced to express luciferase 2 (Luc2) and fluorescent reporter protein tdTomato (GL261 Luc2-tdTomato, CT-2A Luc2-tdTomato). The lentiviral Luc2-tdTomato construct was a gift from Prof. Bentires (University of Basel, Basel, Switzerland). For lentiviral transduction, tumor cells were plated at 5 x 10^4^ cells per well of a 24-well plate 16h prior to transduction. Media was replaced with 0.5 mL antibiotic-free media containing 8 ug mL^-1^ Polybrene (Sigma-Aldrich). 5 μL of lentivirus suspension was added to the cells and incubated for 6 h at 37°C. Afterwards, transduction medium was replaced with normal growth medium and cells were expanded for 4 d and positively sorted for tdTomato expression. Luciferase expression was checked by using in vitro bioluminescence imaging by adding 100 μL of 15 mg mL^-1^ D-luciferin solution (#LUCNA-1G, Goldbio, USA) to the cells and imaged after 2 min using a Fusion FX system (Vilber, France).

### Stereotactic tumor cell injection

Intracranial tumor engraftment was performed at age 8–12 weeks. Mice were being anesthetized with 2.5% isoflurane in an induction chamber. Anesthesia was maintained at 1.5% isoflurane delivered through a nose adaptor on the stereotactic frame. A midline incision was made and a burr hole was drilled at 1 mm posterior to the Bregma, and 2 mm lateral from the midline to the right. Stereotactic injection of tumor cells was done using a Hamilton 10 μL syringe (#80300, 701N, 26s/2”/2, Hamilton) at a depth of 3mm below the dura surface. A microinjection pump was used to inject 5 x 10^4^ tumor cells in 3 μL PBS + 2% FBS total volume at 1 μL/min injection speed. One minute after injection ended, the needle was slowly retracted to avoid reflux of the cell suspension. The scalp wound was closed with sutures (5-0, Polypropylene suture, #8661H, Ethicon, USA).

### In vivo bioluminescent imaging (BLI)

Mice were images by BLI starting at 7 d post tumor injection. Isoflurane-anesthetized mice were injected intraperitoneally (i.p.) with 150 mg kg^-1^ D-luciferin (#LUCNA-1G, Goldbio, USA) and imaged after 5 min using a Newton 7.0 BLI system (Vilber, France). A region of interest (ROI) was drawn around the head and quantitated on mean luminescence (photon count per pixel area). Mice were imaged weekly and checked for neurological symptoms and weighed daily from day 21 onwards.

### In vivo treatments

Tamoxifen (#T5648-5G, Sigma-Aldrich, USA) was dissolved in corn oil (#C8267, Sigma-Aldrich, USA) and 2.5 mg were administered i.p. per mouse and injection, starting from d 7 post tumor injection daily until d 21 and from d 21 onwards, every other day throughout the duration of the experiment. Cre^+^ and Cre^-^ mice were treated with Tamoxifen.

Anti-mouse Ccl2 (clone 2H5, BioXcell) and isotype control (Armenian hamster IgG, BioXcell) were administered at 1 mg kg^-1^ i.p. 3 times per week, starting 7 d after tumor cell injection throughout the duration of the experiment.

Anti-mouse CD47 (clone MIAP410, BioXcell) and isotype control (mouse IgG1, clone MOPC-21, BioXcell) were administered at 4 mg kg^-1^ i.p. on d 11 post tumor injection and at 8 mg kg^-1^ i.p. on d 14, 17 and 20 post tumor injection.

Anti-mouse PD-1 (clone RMP1-14, BioXcell) and isotype control (rat IgG2a, clone 2A3, BioXcell) were administered at 10 mg kg^-1^ i.p. on d 9, 12, 15 and 18 post tumor injection.

Anti-mouse CD8a (clone 53-6.7, BioXcell) for in vivo depletion of CD8^+^ T cells, was administered at 10 mg kg^-1^ i.p. on d −2 and 0 relative to the time of tumor cell injection and depletion was maintained by weekly treatments throughout the duration of the experiment.

### Mouse tumor dissociation

Mice were euthanized by CO_2_-suffocation at endpoint or indicated time points and tumor-bearing cerebral hemisphere without cerebellum was harvested into ice-cold HBSS. On ice, brain tissue was manually minced using razor blades and enzymatically dissociated at 37°C for 30 minutes using the same dissociation buffer as described above. The suspension was filtered through a 70 μm strainer and subjected to a density gradient centrifugation using debris removal solution (#130-109-398, Miltenyi Biotec, Germany) according to manufacturer’s protocol to remove myelin and cell debris. Following ACK-lysis, the remaining myelin- and erythrocytes-depleted tumor cell suspension was used in downstream applications.

### Flow cytometry

Primary mouse and human cells were isolated as described above and stained with Zombie fixable viability dye for 20 min at room temperature (#423102, BioLegend, USA) and pre-incubated with anti-mouse Fc block at 10 μg mL^-1^ (#101320, BioLegend, USA) or anti-human Fc block (#422302, BioLegend, USA). Surface markers were stained with appropriate antibodies for 20 min at 4°C. For intracellular staining, cells were fixed and permeabilized using Cyto-Fast Fix/Perm Buffer Set (#426803, BioLegend, USA), prior to staining for 20 min at room temperature in the dark. For transcription factor staining, cells were fixed, permeabilized and washed using True-Nuclear Transcription Factor Buffer Set (#424401, BioLegend, USA) according to the manufacturer’s protocol. Followed by staining with intracellular antibodies for 30 min at room temperature in the dark. After respective staining protocol, cells were washed twice and resuspended in FACS buffer. For FC, either a Fortessa LSR II (BD Bioscience) or Cytoflex S (Beckman Coulter) flow cytometer were used for cell acquisition and FlowJo software (v.10.8.1, TreeStar) for data analyzation. Cell sorting was performed using a BD FACSAria III or BD FACSMelody (BD Bioscience). We performed compensation using Ultracomp ebeads Compensation Beads (#01-2222-42, Invitrogen, USA), which were stained with appropriate antibody and analyzed on the same voltage and settings. Gates were drawn by using Fluorescent Minus One (FMO) controls. The antibodies used in flow cytometry and cell sorting can be found in Supplementary Tables 3 and 4. Gating strategies are shown within the figures or summarized in Supplementary Data Fig. 1d, e. Either percentage of cell population of interest or Median Fluorescent Intensity (MFI) are reported.

### Imaging flow cytometry

Tumor cell suspensions from *Siglece^fl/fl^* and *Siglece^fl/fl^* x *Sall1^CreERT2^* mice were generated, Fc-receptor blocked and stained with anti-mouse CD11b (clone M1/70, BV650, BioLegend) and anti-mouse CD45 (clone 30-F11, FITC, BioLegend) as described above to identify MG as CD45^low^CD11b^+^ and MdCs as CD45^high^CD11b^+^. DAPI staining was used to exclude dead cells. Imaging flow cytometry was performed on Amnis ImageStreamX Mark II (Merck Millipore). Phagocytosis was analyzed using the IDEAS software onboard internalization-identification algorithm (Amnis), which identifies tdTomato signal within MG events. Pictures were taken at 40X magnification at low speed, high sensitivity mode. Full gating strategy is shown in Supplementary Data Figure 1h.

### scRNA-seq analysis of immune cells

Tumors from *Siglece^fl/fl^* and *Siglece^fl/fl^* x *Sall1^CreERT2^* mice were dissociated as described above and cells were stained with Zombie fixable viability dye, Fc-receptor blocked and stained with CD45 and CD11b. Live, CD45^+^ and/or CD11b^+^ immune cells were sorted into ice-cold PBS with 0.4% BSA. Cells were kept on ice and brought to the Genomics Facility Basel at the Department of Biosystems Science and Engineering of the ETH Zurich, Basel. Single cell capture and cDNA and library preparation were performed with Chromium Next GEM Single Cell 5’ Reagent Kits v2 (10x Genomics, CA, USA) following manufacturer’s protocol with the goal of loading 10,000 cells per sample. After library preparation, libraries were sequenced using the NovaSeq 6000 systems (Illumina, CA, USA) to produce paired-end 91nt R2 reads with a targeted minimum sequencing depth of 20,000 genes per cell.

The dataset was analyzed by the Bioinformatics Core Facility, Department of Biomedicine, University of Basel. Read quality was assessed with the FastQC tool (version 0.11.5). Sequencing files were processed with STARsolo (STAR version 2.7.10a) [65] to perform sample and cell demultiplexing, and alignment of reads to the mouse genome (mm10; supplemented with the Cre, tdTomato eGFP construct sequences) and UMI counting on gene models from Ensembl 102. The options “*--outFilterType=BySJout -- outFilterMultimapNmax=10 --outSAMmultNmax=1 --outFilterScoreMin=30 -- soloCBmatchWLtype=1MM_multi_Nbase_pseudocounts --soloUMIlen=10 -- soloUMIfiltering=MultiGeneUMI_CR --soloUMIdedup=1MM_CR --soloCellFilter=None -- clipAdapterType=False --soloType=CB_UMI_Simple –soloStrand=Reverse – soloMultiMappers=EM*” were used for STARsolo.

For each sample, empty droplets were detected and removed using the *emptyDrops()* function from the Bioconductor DropletUtils package (version 1.14.2; using 5000 iterations, the option *test.ambient=TRUE*, a lower threshold of 100 UMIs and an FDR threshold of 0.1%) [66]. Processing of the UMI counts matrix was performed using the Bioconductor packages scran (version 1.24.0) [67, 68] and scatter (version 1.24.0) [69], following mostly the steps illustrated in the OSCA book (http://bioconductor.org/books/3.15/OSCA/) [68, 70]. A first step of filtering for low-quality cells was done based on library size (at least 600 UMI counts per cell), the number of detected genes (at least 400 genes detected) and the percentage of reads mapping to mitochondrial genes (larger than 0% and lower than 20%), based on the distribution observed across cells. The presence of doublet cells was investigated with the scDblFinder package (version 1.10.0), and suspicious cells were filtered out (score > 0.9). UMI counts were normalized with size factors estimated from pools of cells created with the scran package *quickCluster()* function [67, 71]. To distinguish between genuine biological variability and technical noise, we modeled the variance of the log-expression across genes using a Poisson-based mean-variance trend. The scran package *denoisePCA()* function was used to denoise log-expression data by removing principal components corresponding to technical noise. Cell cycle phase was inferred using the markers provided in the package Seurat (version 4.1.1) and the *CellCycleScoring()* function. Systematic differences between samples were removed using the *fastMNN()* function (d=50, k=20) of the batchelor package (version 1.12.3) [72], run on the most hypervariable genes across cells (excluding MT genes and ribosomal protein genes, as well as genes at high level in ambient RNA).

Clustering of cells was performed with hierarchical clustering on the Euclidean distances between cells (with Ward’s criterion to minimize the total variance within each cluster [73]; package cluster version 2.1.3). The number of clusters used for following analyses was identified by applying a dynamic tree cut (package dynamicTreeCut, version 1.63-1) [74], and the clustering resolution was adapted using the argument deepSplit (default 1). The *scoreMarkers()* function of the scran package was used to find the best markers across clusters. Based on their high levels of MT reads, low number of detected genes and lack of specific cell-type markers, some clusters were deemed composed of low-quality cells and filtered out.

The best markers across clusters and a list of known cell type-specific genes were used for cell type annotation. In addition, the Bioconductor package SingleR (version 1.10.0) was used for unbiased cell-type annotation of the cells [75] using as reference datasets (i) sorted bulk microarray and RNA-seq samples from the ImmGen project [76], (ii) a collection of mouse RNA-seq samples annotated to 28 cell types, available through the celldex Bioconductor package; (iii) a collection of scRNA-seq datasets from virus-specific CD8 T cells, tumor-infiltrating T cells, and virus-specific CD4 T cells available at https://spica.unil.ch/refs [77, 78]. A microglia and a macrophage signature scores were defined by taking to 1-to-1 orthologs in mouse (Using Ensembl Compara [79]) of genes from the lists obtained in [80], and averaging their center and scaled expression levels. To improve the annotation of the T cells and monocytes/macrophages/microglia clusters, a reclustering was performed based on the set of most hypervariable genes within cells of each of these subsets. Finally, the SingleR high-quality assignments (pruned scores) and the signature scores were used to manually derive a consensus cell type annotation for each cluster.

After quality filtering, the resulting dataset consisted of UMI counts for 28,644 cells, ranging from 1,410 to 5,856 cells per sample. A two-dimensional t-distributed stochastic neighbour embedding (tSNE) used for visualization of cells was calculated using the batch-corrected matrix of low-dimensional coordinates for each cell (perplexity of 30).

Differential abundance analysis across cell types between *Siglece^fl/fl^* x *Sall1^CreERT2^* and *Siglece^fl/fl^* conditions was performed using the *limma-voom* method [81, 82]. Differential abundance of cell types was considered to be significant at a false discovery rate (FDR) lower than 5 %.

Differential expression between conditions, stratified by annotated cell type, was performed using a pseudo-bulk approach, summing the UMI counts of cells from each cell type in each sample when at least 20 cells could be aggregated. The aggregated samples were then treated as bulk RNA-seq samples [83] and for each pairwise comparison genes were filtered to keep genes detected in at least 5% of the cells aggregated. Because the number of cells in the microglia clusters was too low, we pooled both clusters (‘MG_1’ and ‘MG_2’) together for the pseudo-bulk analysis. The package edgeR (version 3.38.4) [84] was used to perform TMM normalization [85] and to test for differential expression with the Generalized Linear Model (GLM) framework. Genes with a false discovery rate (FDR) lower than 5% were considered differentially expressed. Gene set enrichment analysis was performed with the function camera [86] on gene sets from the Molecular Signature Database (MSigDB, version 7.5.1) [36, 87]. We retained only sets containing more than 5 genes, and gene sets with a FDR lower than 5% were considered as significant.

### Antigen cross-presentation assay

5 x 10^4^ CT-2A Luc2-tdTomato cells were intracranially injected into *Siglece^fl/fl^* and *Siglece^fl/fl^* x *Cx3cr1^CreERT2^* mice and tamoxifen i.p. injected on days 7 to 12 after tumor injection. On d 15, tumor-bearing hemispheres without cerebellum were harvested and dissociated as described above. CD11b^+^ cells were then isolated using the CD11b+ microbeads (#130-093-636, Miltenyi) according to the manufacturer’s instructions on an AutoMACS (Miltenyi). Isolated cells were plated at 5 x 10^4^ cells per well of a tissue-culture treated 96-well flat-bottom plate (Costar) in 100 μL IMDM medium (Gibco) with 10% FBS. Full-length ovalbumin protein (#vac-pova, InvivoGen, USA) was added at 100 μg mL^-1^ to the cells and incubated for 20 h at 37°C. Cells were collected with TrypLE (Gibco), Fc-receptor blocked and stained with anti-mouse CD11b (clone M1/70, FITC, BioLegend), anti-mouse CD45 (clone 30-F11, BV605, BioLegend) and APC-labeled anti-SIINFEKL/H-2Kb (#17-5743-80, eBioscience, USA), which specifically detects ovalbumin-derived SIINFEKL peptide bound to H-2Kb. DAPI nuclear stain was used to exclude dead cells. Antigen cross-presentation was assessed using a Cytoflex S flow cytometer (Beckman Coulter) and was measured as the percentage of APC^+^ cells within CD45^high^CD11b^+^ MdCs and CD45^low^CD11b^+^ MG. Differences in antigen cross-presentation were reported as fold change between cells derived from *Siglece^fl/fl^* x *Cx3cr1^CreERT2^* and *Siglece^fl/fl^* mice.

### OT-I CD8^+^ T cell isolation

Spleens from Rag2/OT-I transgenic mice were carefully removed and washed with PBS containing 2% FBS under sterile conditions. Tissue was mechanically dissociated by crushing it with a syringe piston in a 10 cm dish. Cell suspension was passed through a 70 μm strainer and washed. CD8^+^ T cell isolation was performed using EasySep Mouse CD8^+^ T cell Isolation Kit (#19853, STEMCELL Technologies, USA) according to manufacturer’s protocol.

### T cell cross-priming assay

CD11b^+^ cells were isolated, plated and treated as described in antigen cross-presentation assay. After 20 h incubation of the CD11b^+^ cells, naive OT-I CD8^+^ T cells were isolated and added to the CD11b^+^ cells at 2 x 10^5^ cells per well and incubated for an additional 24 h. For T cell activation analysis, cells were collected and stained with CD8a (clone 53-6.7, FITC), CD25 (clone PC61, APC), CD69 (clone H1.2F3, PerCP/Cy5.5) and CD44 (clone IM7, PE) (all from BioLegend) and acquired using a Cytoflex S flow cytometer (Beckman Coulter). Of note, at two weeks post tumor injection (timepoint for CD11b^+^ cells isolation), we observed a reduced proportion of infiltrating MdCs among CD11b^+^ cells in *Siglece^fl/fl^* x *Cx3cr1^CreERT2^* mice compared to *Siglece^fl/fl^* control, probably due to the delayed tumor growth, since the proportion of MdCs among CD11b^+^ cells showed a high level of correlation with the BLI count (Supplementary Data Fig. 4a). Therefore, to account for the reduced proportion of the cross-presenting MdCs among the isolated CD11b^+^ cells in the *Siglece^fl/fl^* x *Cx3cr1^CreERT2^* group, T cell activation marker expression was normalized to the mean bioluminescence count.

### Proteomics analysis of MdCs

1.5 x 10^5^ MdCs per tumor-bearing mouse brain were sorted and thoroughly washed using protein-free PBS, pelleted by centrifugation and the cell pellet was snap frozen on dry ice and kept in −80°C. For mass spectrometry-based proteomics, frozen cell pellets were brought to the Proteomics Core Facility of the University of Basel. There, cells were lysed with 50 µL of lysis buffer (1% Sodium deoxycholate (SDC), 10 mM Tris(2-carboxyethyl)phosphine (TCEP), 100 mM Tris, pH=8.5) using 10 cycles of sonication (30 s on, 30 s off per cycle) on a Bioruptor (Dianode). Following sonication, proteins were reduced by TCEP at 95° C for 10 min. Proteins were alkylated using 15 mM chloroacetamide at 37° C for 30 min and further digested using sequencing-grade modified trypsin (1/50 w/w, ratio trypsin/protein; Promega, USA) at 37° C for 12 hours. After digestion, the samples were acidified using TFA (final 1%). Peptide desalting was performed using iST cartridges (PreOmics, Germany) following the manufacturer’s instructions. After drying the samples under vacuum, peptides were stored at −20° C

Dried peptides were resuspended in 0.1% aqueous formic acid and subjected to LC– MS/MS analysis using an Orbitrap Eclipse Tribrid Mass Spectrometer fitted with Ultimate 3000 nano system and a FAIMS Pro interface (all Thermo Fisher Scientific) and a custom-made column heater set to 60°C. Peptides were resolved using a RP-HPLC column (75μm × 30cm) packed in-house with C18 resin (ReproSil-Pur C18–AQ, 1.9 μm resin; Dr. Maisch GmbH) at a flow rate of 0.3 μL/min. The following gradient was used for peptide separation: from 2% B to 12% B over 5 min to 30% B over 70 min to 50% B over 15 min to 95% B over 2 min followed by 18 min at 95% B then back to 2% B over 2 min followed by 8 min at 2% B. Buffer A was 0.1% formic acid in water and buffer B was 80% acetonitrile, 0.1% formic acid in water.

The mass spectrometer was operated in DDA mode with a cycle time of 3 seconds between master scans. Throughout each acquisition, the FAIMS Pro interface switched between CVs of −40 V and −70 V with cycle times of 1.5 s and 1.5 s, respectively. MS1 spectra were acquired in the Orbitrap at a resolution of 120,000 and a scan range of 375 to 1500 m z^-1^, AGC target set to 10^6^ and maximum injection time set to 50 ms. Precursors were filtered with precursor selection range set to 375–1500 m z^-1^, monoisotopic peak determination set to “Peptide”, charge state set to 2 to 5, a dynamic exclusion of 30 s.

Precursors selected for MS2 analysis were isolated in the quadrupole with a 1.4 m z^-1^ window and collected for a maximum injection time of 35 ms with AGC target set to “Standard”. Fragmentation was performed with a CID collision energy of 30% and MS2 spectra were acquired in the IT at scan rate “Turbo”.

The raw files were using FragPipe v17.1 software with standard LFQ settings. In brief, the spectra were searched against a mouse protein database (retrieved from Uniprot on 21.04.2022) and commonly observed contaminants by the MSFragger v3.4 search engine [88] using the following criteria: full tryptic specificity was required, 2 missed cleavages were allowed, carbamidomethylation (C) was set as fixed modification, oxidation (M) and acetylation (Protein N-term) were applied as variable modification, minimum and maximum peptide lengths were set to 7 and 50 respectively. LFQ quantification was performed with IonQuant [89] with match between runs (MBR) algorithm was enabled. The database search results were filtered to a false discovery rate (FDR) of 1 %. Differential quantitative analysis was performed using MSstats package (v4.0.1) [90]. The KEGG pathway over-representation analysis was performed using the ClusterProfiler package in R. The genes of interest were selected according to log2 fold change, we selected genes with values between −1 and 1 (n=104) the universe comprised all the genes from the dataset, minGSSize = 3, maxGSSize = 800, pAdjustMethod = “fdr”, pvalueCutoff = 1 and qvalueCutoff = 1.

### RNA extraction and quantitative real-time PCR (qPCR)

Total RNA was isolated form FACS-sorted MdCs using the Direct-zol RNA MiniPrep plus kit (#R2072, Zymo Research, USA). mRNA was transcribed with iScript cDNA Synthesis Kit (#1708891, Bio-Rad, USA) and qPCR was performed on a CFX96 Real-Time System (Bio-Rad) using SYBR Green qPCR Mastermix (#330500, Qiagen, Germany). For NF-κB target gene analysis, primer array of the RT^2^ Profiler PCR Array (#PAMM-025ZD, Qiagen, Germany) was used according to manufacturer’s protocol. Additionally, QuantiTect Primer Assay for *Siglece* (#QT00135597, Qiagen, Germany) was used. *Actb, B2m, Gapdh, Gusb* and *Hsp90ab1* were used to normalize signal expression. Fold regulation comparison were performed between control and treatment samples using the 2^-1′1′Ct^ method [91].

### Staining for Siglec-9 and Siglec-E ligands on tumor cells

For flow cytometric-based analysis of Siglec ligands, single cell suspensions of tumor cells were blocked, live/dead stained and incubated with recombinant human Siglec-9 Fc chimera (#1139-SL-050) or recombinant mouse Siglec-E Fc chimera (#5806-SL-050) both from R&D systems, USA for 20 min on ice. For detection, anti-human IgG (clone HP6017, APC) or anti-mouse IgG2a (clone RMG2a-62, APC or BV421) both from BioLegend, were incubated for 20 min on ice. Samples were acquired on a CytoFLEX S flow cytometer (Beckman Coulter).

### Generation of CT-2A^ΔGNE^ tumor cells

Knockout of *Gne* in CT-2A tumor cells was performed using CRISPR/Cas9 mediated gene editing. The plasmid containing the guide RNAs were kindly provided by Dr. Stanczak (University Hospital Basel, Basel, Switzerland) [92]. Tumor cells were transiently transfected with the plasmid using FuGENE HD transfection reagent (#E2311, Promega, USA) according to manufacturer’s protocol and GFP^+^ cells were sorted. After their recovery and expansion, cell surface sialylation was analyzed by flow cytometry using recombinant Siglec-E Fc chimera as described above. For enzymatic desialylation control, cells were pre-incubated with 20 mU ml^-1^ Vibrio cholerae-derived neuraminidase (sialidase) (#11080725001, Roche, Switzerland) for 1 h at 37°C on a cell shaker. The wildtype parental cell line, as well as transfected and sorted cells showing normal cell surface sialylation were used as controls. CT-2A^ΔGNE^ cells were compared to CT-2A^WT^ with regard to their in vitro viability and proliferation, as well as their in vivo tumorigenicity in wildtype C57BL/6 mice. Proliferation was assessed by a MTT assay (#M2128, Sigma-Aldrich, USA) and viability of cells was measured by DAPI-stain exclusion.

### Ex vivo perfusion bioreactor

Bioreactor cultures under perfusion were performed according to the manufacturer’s instructions (Cellec Biotek AG, Basel, Switzerland) and as previously described [47]. In brief, fresh, intact GBM tissue explants were placed into ice-cold Neurobasal-A medium (#21103049, Life Technologies, USA) and immediately taken to the laboratory (less than 30 min). Tumor tissue was cut into ∼20 to 30 mm^3^ fragments. Silicone adaptors and ethylene-tetrafluoroethylene copolymer mesh grids were arranged on the top and bottom of the tissue, and placed into U-CUP perfusion chambers (Cellec Biotek AG). The perfusion media consisted of a 50:50 mix of Neurobasal-A medium (#21103049, Life Technologies, USA) and Dulbecco’s modified Eagle’s medium/F12 medium (#21331020, Gibco, USA) supplemented with non-essential amino acids (1×; #M7145, Sigma-Aldrich, USA), 1 mM sodium pyruvate (#S8636, Sigma-Aldrich, USA), 44 mM sodium bicarbonate (#25080060, Gibco, USA), 25 mM Hepes (#156301, Gibco, USA), 4 mM l-alanyl-l-glutamine (#25-015-CI, Corning, USA), antibiotic-antimycotic (1×; #15240062, Gibco, USA), human recombinant epidermal growth factor (20 ng/ml; #236-EG-01M, R&D systems, USA), human recombinant fibroblast growth factor (20 ng/ml; #100-18B, PeproTech, UK), heparin sulfate (10 ng/ml; 100-18B, STEMCELL Technologies, USA), and 5% human serum (#H4522-100ML, Sigma-Aldrich, USA). The perfusion flow rate was set at 0.47 ml/min, resulting in a superficial flow velocity of 100 μm/s. Starting from day 0, bioreactor media were either left untreated or supplemented with anti– Siglec-9 antibody (5 μg mL^-1^; clone 191240, #MAB1139-500, R&D systems, USA). On day 5, culture media were frozen at −80°C and tissues were fixed in 10% neutral buffered formalin (#HT501128, Sigma-Aldrich, USA) for 24 h, afterwards transferred to 70% ethanol and embedded in paraffin.

### Multiplexed secreted protein analysis in explant culture media

After five days of culture, we measured soluble proteins including cytokines and chemokines in the media of each bioreactor using the proximity extension assay (PEA) technology developed by Olink Proteomics. Bioreactor media were centrifuged to remove cell debris, and 92 secreted proteins, including cytokines, chemokines, and soluble cell membrane proteins, were measured externally by PEA technology using the Olink Target 96 Immuno-Oncology panel (Olink, Sweden). Differences in normalized protein expression (NPX), Olink’s arbitrary unit on a log_2_ scale, were reported as fold change to the respective control.

### Pharmacoscopy

Tumor dissociation was performed as described above and dissociated single cells were resuspended in DMEM media supplemented with 10% FBS, 25 mM HEPES and 1% Pen/Strep and were seeded at 10^4^ cells/well (in 50 μl/well) into clear-bottom, tissue-culture treated, CellCarrier-384 Ultra Microplates (Perkin Elmer, Waltham, Massachusetts, USA). Anti– Siglec-9 antibody (5 μg mL^-1^; clone 191240, #MAB1139-500, R&D systems, USA) or human IgG isotype control (#31154, Thermo Fisher Scientific) was added to the single cell suspensions and cultured at 37°C, 5% CO_2_ for 48 h. Per patient there were three technical replicate wells for the anti-Siglec-9 treatment and six technical replicate wells for the IgG control. Subsequently, cells were fixed with 4% PFA (Sigma-Aldrich), blocked with PBS containing 5% FBS, 0.1% TritonX and DAPI (4 ug/mL, #422801, Biolegend) for one hour at RT and stained with the following antibodies: Alexa Fluor 488 anti-NESTIN (#656812, Biolegend), Alexa Fluor 555 anti-S100B (#ab274881, Abcam), Alexa Fluor 647 anti-CD45 (#368538, Biolegend) overnight at 4°C. Imaging of the 384 well plates was performed with an Opera Phenix automated spinning-disk confocal microscope at 20x magnification (Perkin Elmer). Single cells were segmented based on their nuclei (DAPI channel) using CellProfiler 2.2.0. Downstream image analysis was performed with MATLAB R2021b. Marker positive cell counts for each condition were derived based on a linear threshold of the histograms of each channel, were averaged across each well and compared between aSiglec-9 treatment and the IgG control group.

### Statistical analysis

scRNA-seq and proteomic statistical analysis was completed as described above. All other statistical analyses were performed using GraphPad Prism (GraphPad Software v.9.4.0). Numbers of experimental replicates, numbers of independent experiments and statistical tests used are given in the figure legends. In general, when two groups were compared, significance was determined using an unpaired two-tailed Student’s t test. For comparing more than two groups, one-way ANOVA was applied and for comparing a quantitative variable between two groups, two-way ANOVA was used. Survival data were analyzed using the log-rank Mantel-Cox test or restricted mean survival time (RMST). RMST analysis was used to account for the presence of censoring. The calculations were performed in R using the survRM2 package. We used max τ (largest observed time in each of the two groups) and the differences in RMST between subgroups were calculated as 95% CIs with P values. *P* values < 0.05 were considered as statistically significant. No statistical methods were used to predetermine sample size. Data collection and analysis were not performed blind to the conditions of the experiments. Outliers were removed using GraphPad Outlier Calculator (https://www.graphpad.com/quickcalcs/Grubbs1.cfm).

### Graphical illustrations

All graphical illustrations were created with <u>BioRender.com</u>.

## Data availability

The scRNA-seq dataset is available on GEO under accession number <u>GSE212616</u>.

## Autor contributions

G.H., H.L. and P.S., conceived and planned the project. P.S. designed, performed and analyzed most experiments and wrote the manuscript. J.R. and S.H. provided bioinformatical support and performed computational analysis. A.B., N.T., J.W., S.L., B.S., T.A.M., M.-F.R., T.S., D.K. and M.M performed and analyzed experiments. M.W., T.W. supervised and coordinated experiments. G.H. and H.L. supervised and coordinated the study and critically revised the manuscript. All authors reviewed the paper and approved its final version.

## Acknowledgments

We are grateful to the patients and their families for their consent to donate tissue to our brain tumor biobank. Calculations were performed at sciCORE (http://scicore.unibas.ch/) scientific computing center at the University of Basel. This work was supported by a Swiss Cancer Research MD-PhD Grant (MD-PhD-4818-06-2019) to P.S. as well an Alumni Medizin Basel grant to P.S.; Swiss National Science Foundation Professorial Fellowship (PP00P3_176974); the ProPatient Forschungsstiftung, University Hospital Basel (Annemarie Karrasch Award 2019); Swiss Cancer Research Grant (KFS-4382-02-2018) to G.H.; the Department of Surgery, University Hospital Basel, to G.H. and P.S.; and by The Brain Tumour Charity Foundation, London, UK (GN-000562) to G.H. We thank Prof. Alfred Zippelius, Prof. Jan Niess and Prof. Ryuichi Nishinakamura for providing genetic mouse lines; Dr. Manuele Muraro for technical help with the perfusion bioreactors; Dr. Claudio Giachino and Dr. Elena Parmigiani from Prof. Verdon Taylor lab for providing the PDGF^+^*Trp53^-^* mouse glioma cell line; Dr. Michal Stanczak for providing the GNE-KO plasmid; Dr. Anne Bärenwaldt and Dr. Natalia Rodrigues Mantuano for discussions and advice; and the genomics, proteomics, flow cytometry and animal core facilities of the University of Basel, Switzerland for technical and logistical support.

## Supplementary Data Figures

**Supplementary Data Fig. 1.**
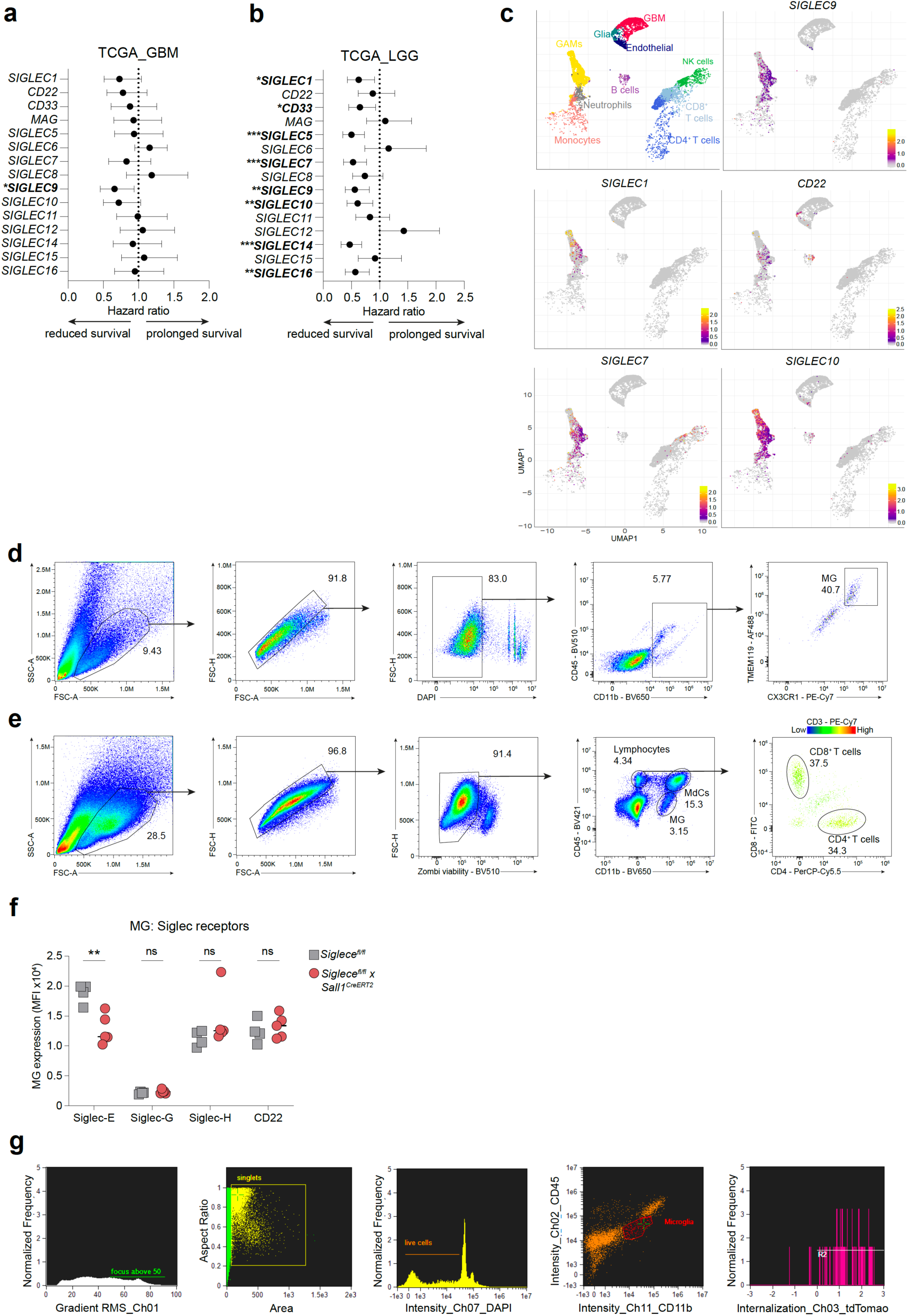
High *SIGLEC9* expression significantly correlates with reduced survival in GBM patients and represents a novel immunotherapeutic target. **a**, **b**, Forest plots showing hazard ratios for overall survival associated with high expression of human Siglecs using The Cancer Genome Atlas (TCGA) RNA-seq dataset of human GBM (**a**) and low-grade glioma (LGG) (**b**) patients. The median mRNA expression value was selected as cutoff for high and low expression groups. A Cox proportional hazard analysis was performed using the GlioVis data portal [22] and hazard ratios with 95% confidence intervals are shown. **c**, Annotated UMAP plot showing scRNA-seq analysis of five primary human GBM tumors [23] (upper left) and UMAP plots showing expression pattern of selected Siglecs which exhibit a hazard ratio for overall survival of < 0.85 in GBM patients, as shown in Supplementary Data Fig. 1a. **d**, Gating strategy to identify human GBM-associated MG. **e**, Gating strategy to identify mouse glioma-associated immune cells. **f**, Flow cytometry analysis of selected mouse Siglecs on MG from *Siglece^fl/fl^* and *Siglece^fl/fl^* x *Sall1^CreERT2^* mice bearing CT-2A tumors (n = 4-5 mice per group). Experiment was performed once. **g**, Gating strategy for imaging flow cytometry to identify MG tumor cell phagocytosis using the IDEAS software onboard internalization-identification algorithm. Statistics: Data are shown as median (**f**), two-way ANOVA with Sidak’s corrected multiple comparison test (**f**). \**p* ≤ 0.05, \*\**p* ≤ 0.01, \*\*\**p* ≤ 0.001.

**Supplementary Data Fig. 2.**
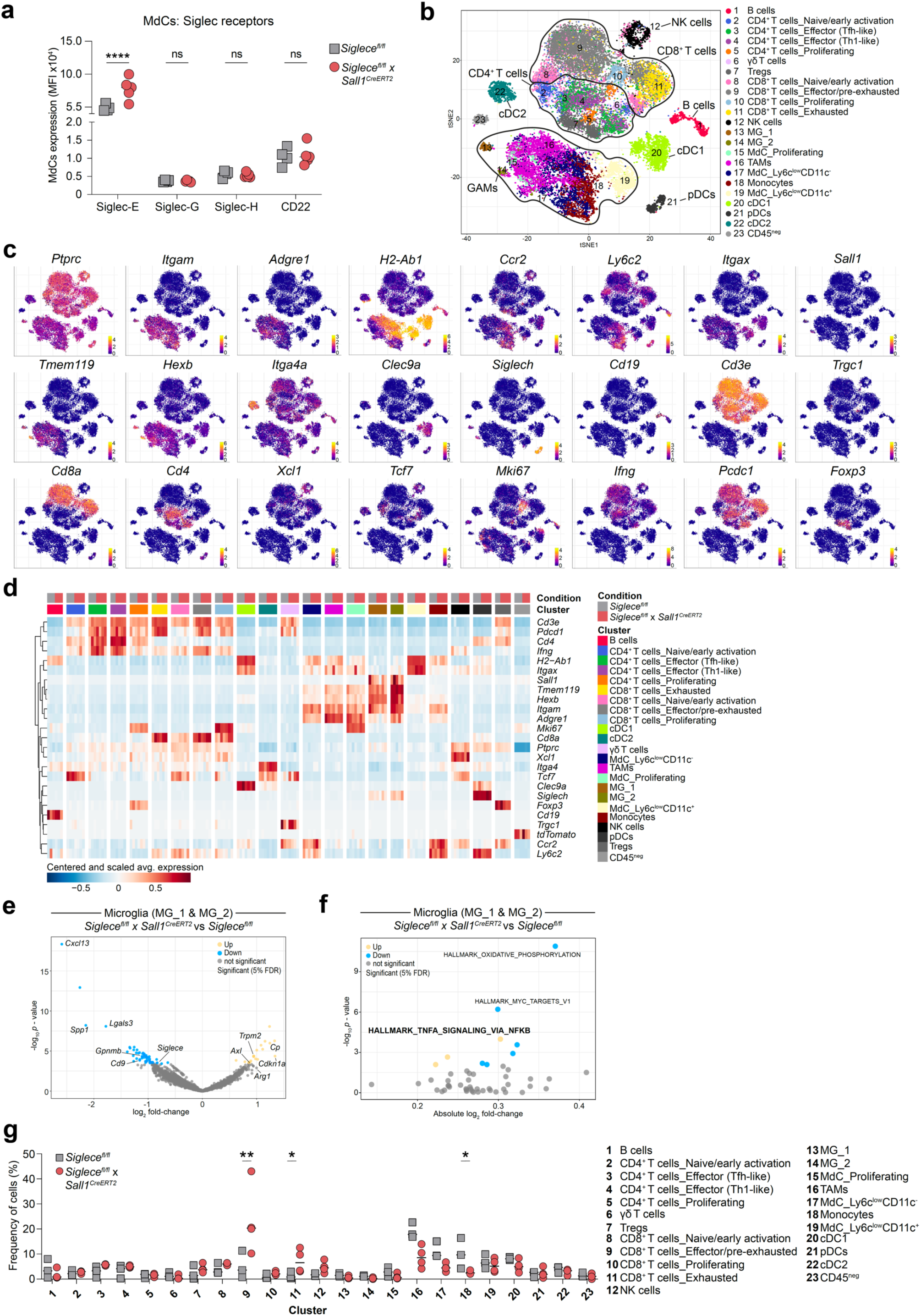
scRNA-seq analysis of immune cells from *Siglece^fl/fl^* and *Siglece^fl/fl^* x *Sall1^CreERT2^* mice. **a**, Flow cytometry analysis of selected mouse Siglecs on MdCs from *Siglece^fl/fl^* and *Siglece^fl/fl^* x *Sall1^CreERT2^* mice bearing CT-2A tumors (n = 4-5 mice per group). Experiment was performed once. **b**, tSNE plot showing the clustering and annotation of cells from *Siglece^fl/fl^* and *Siglece^fl/fl^* x *Sall1^CreERT2^* mice bearing CT-2A tumors. **c**, tSNE plots showing the expression of selected cell-type specific markers. Expression is shown as normalized log_2_ counts **d**, Heatmap displaying log-normalized expression of selected marker genes, averaged across cells of each cluster and sample, and centered and scaled by row. **e**, Volcano plot showing the differential expression analysis results between *Siglece^fl/fl^* x *Sall1^CreERT2^* and *Siglece^fl/fl^* for microglia from the MG_1 & MG_2 clusters. **f**, Scatter plot showing the results of the corresponding GSEA performed on the MSigDB Hallmark collection. **g**, Relative frequencies of *Siglece^fl/fl^* (gray) and *Siglece^fl/fl^* x Sall1^CreERT2^ (red) cells in each cluster. Symbols represent biological replicates. Statistics: Data are shown as median (**a**, **g**), two-way ANOVA with Sidak’s corrected multiple comparison test (**a**), *limma-voom* method with FDR<5% (**g**). \**p* ≤ 0.05, \*\**p* ≤ 0.01, ****p ≤ 0.0001.

**Supplementary Data Fig. 3.**
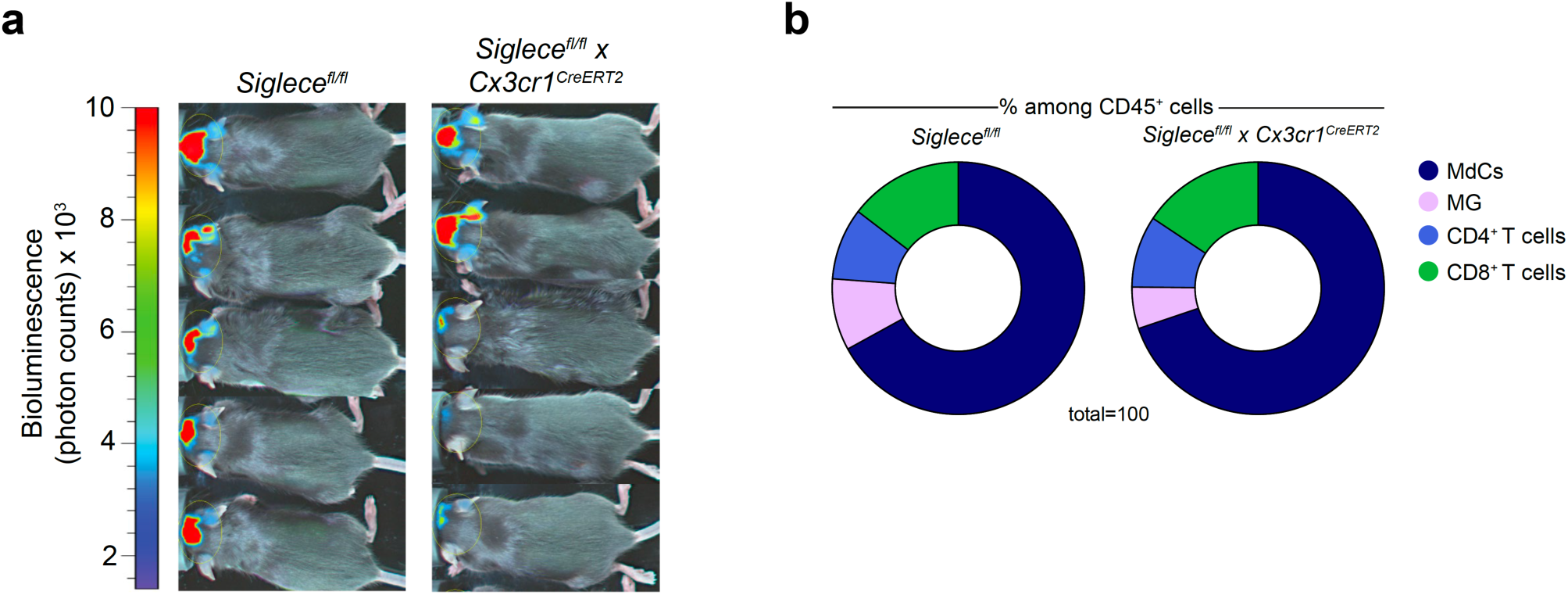
*Siglece^fl/fl^* x *Cx3cr1^CreERT2^* glioma mouse model. **a**, Representative bioluminescence images of day 21 post tumor injection in *Siglece^fl/fl^* and *Siglece^fl/fl^* x *Cx3cr1^CreERT2^* mice engrafted with CT-2A Luc2-tdTomato tumors. Image representative of two independent experiments. **b**, Pie charts represent relative frequencies for main immune cell clusters identified among CD45^+^ cells within *Siglece^fl/fl^* and *Siglece^fl/fl^* x *Cx3cr1^CreERT2^* mice (n = 6 mice per group). Results shown are from one experiment, representative of two independent experiments.

**Supplementary Data Fig. 4.**
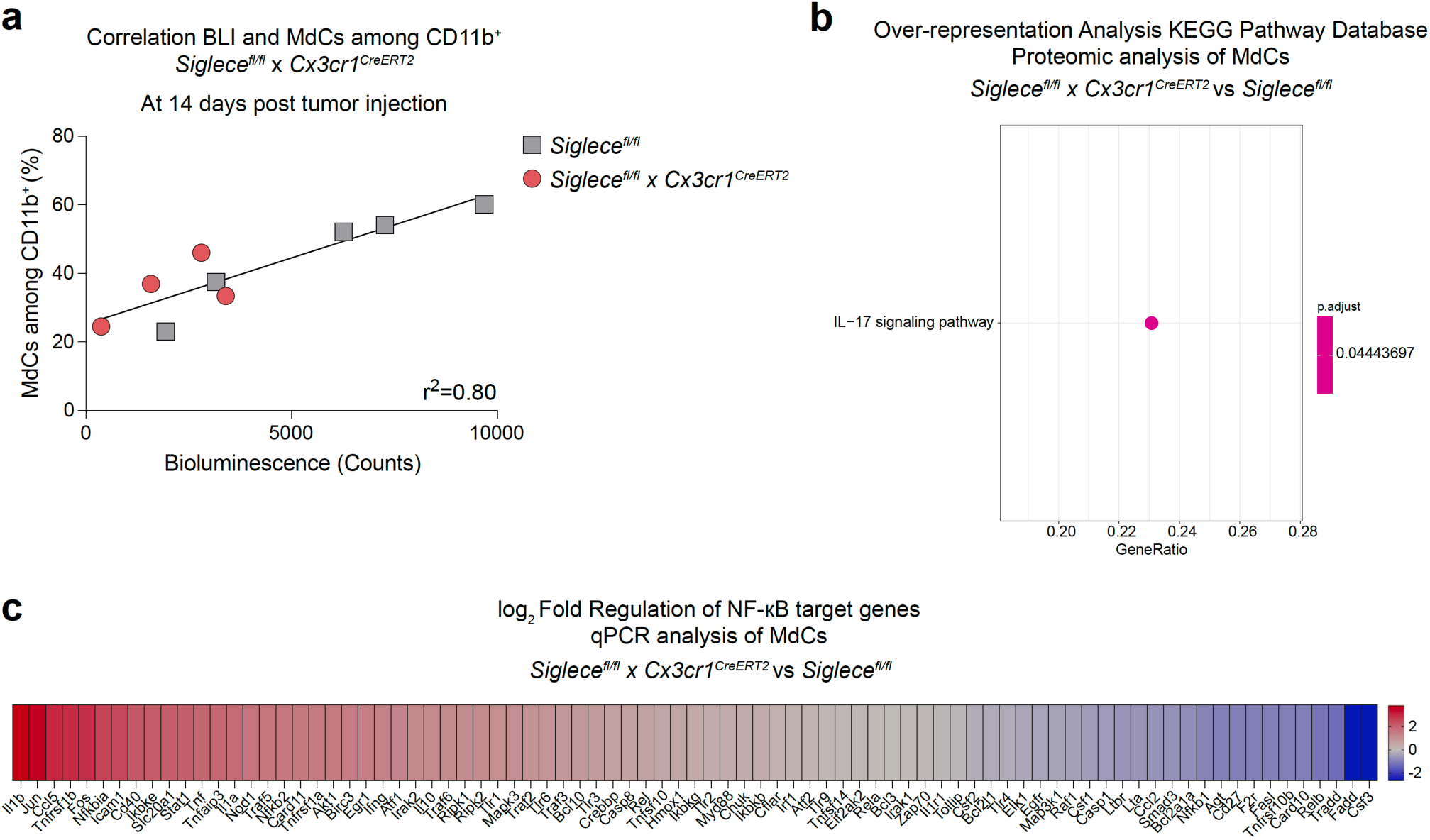
Siglec-E deficient MdCs show increased activity. **a**, Correlation between MdC influx among CD11b^+^ cells and tumor size (measured as bioluminescence count) at day 14 post tumor injection in *Siglece^fl/fl^* and *Siglece^fl/fl^* x *Cx3cr1^CreERT2^* mice (n = 4-5 mice per group). **b**, Over-representation analysis (KEGG Pathway Database) of the proteomic analysis of MdCs sorted from *Siglece^fl/fl^* and *Siglece^fl/fl^* x *Cx3cr1^CreERT2^* mice. **c**, Heat map displaying log_2_ Fold Regulation of NF-κB target genes between MdCs from *Siglece^fl/fl^* and *Siglece^fl/fl^* x *Cx3cr1^CreERT2^* mice. Statistics: Simple linear regression analysis was done to compute r^2^ (**a**), for detailed proteomic statistical analysis, please refer to Methods section.

**Supplementary Data Fig. 5.**
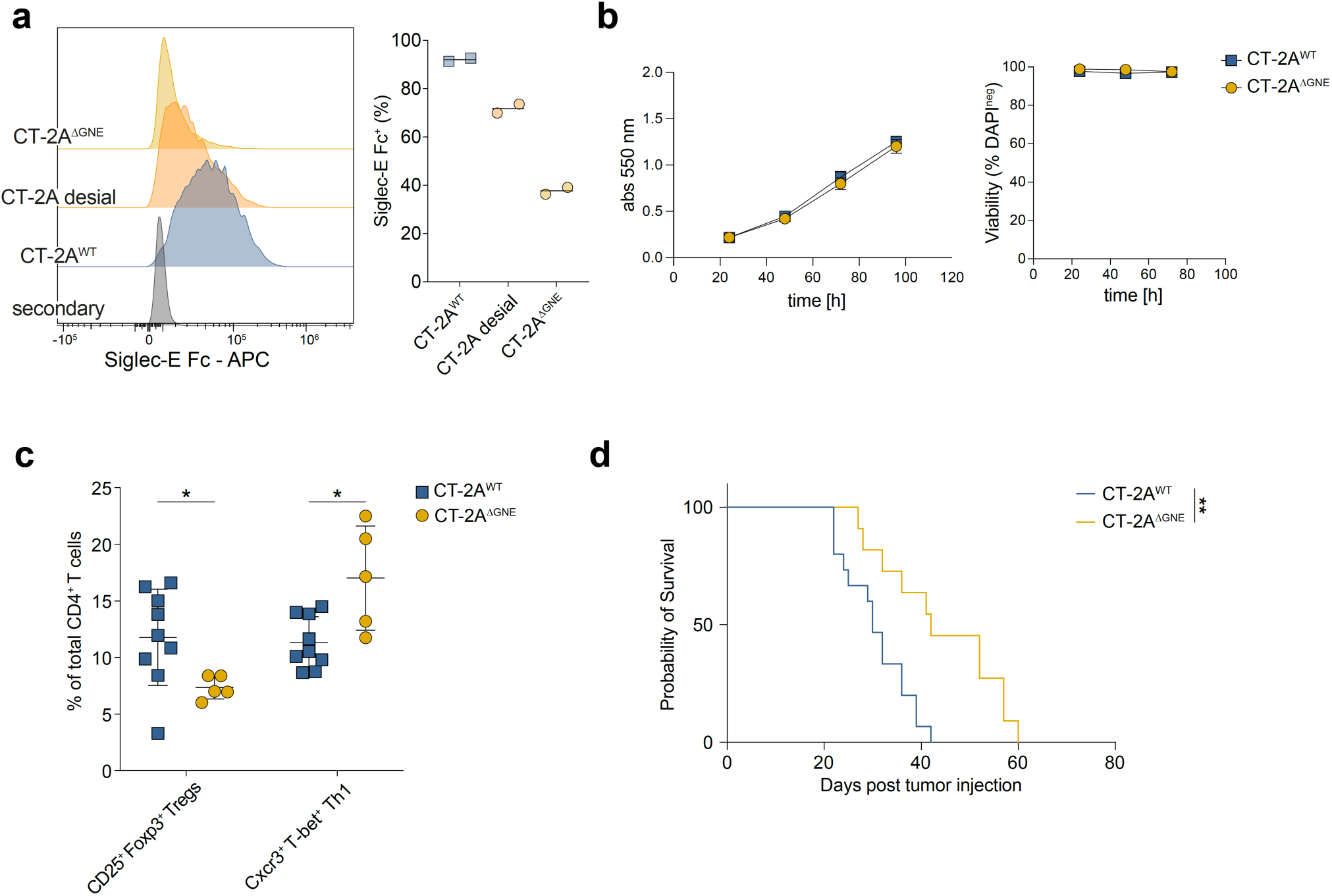
Generation of genetically desialylated CT-2A^ΔGNE^ cells. **a**, Representative histograms (left) and quantification (right) of binding of Siglec-E Fc to CT-2A^WT^, enzymatically desialylated CT-2A cells (CT-2A desial) and GNE-deficient CT-2A cells (CT-2A^ΔGNE^) (n = two independent experimental replicates). **b**, In vitro proliferation (left) and cell viability (right) of CT-2A^ΔGNE^ and CT-2A^WT^ cells (n = two independent experimental replicates). **c**, Flow cytometry analysis showing percentage of CD25^+^ Foxp3^+^ Tregs and Cxcr3^+^ T-bet^+^ Th1 of total CD4^+^ T cells between CT-2A^WT^ and CT-2A^ΔGNE^ injected C57BL/6 wildtype mice (n =5-9 mice per group). Results were pooled from two independent experiments. **d**, Survival of CT-2A^WT^ and CT-2A^ΔGNE^ injected C57BL/6 wildtype mice (n =11-15 mice per group). Results were pooled from two independent experiments. Statistics: Data are presented as median (**a**) and mean ± SD (**b**, **c**), two-way ANOVA with Holm-Sidak’s corrected multiple comparison test (**c**), log-rank Mantel-Cox test (**d**). *p ≤ 0.05, **p ≤ 0.01.

**Supplementary Data Fig. 6.**
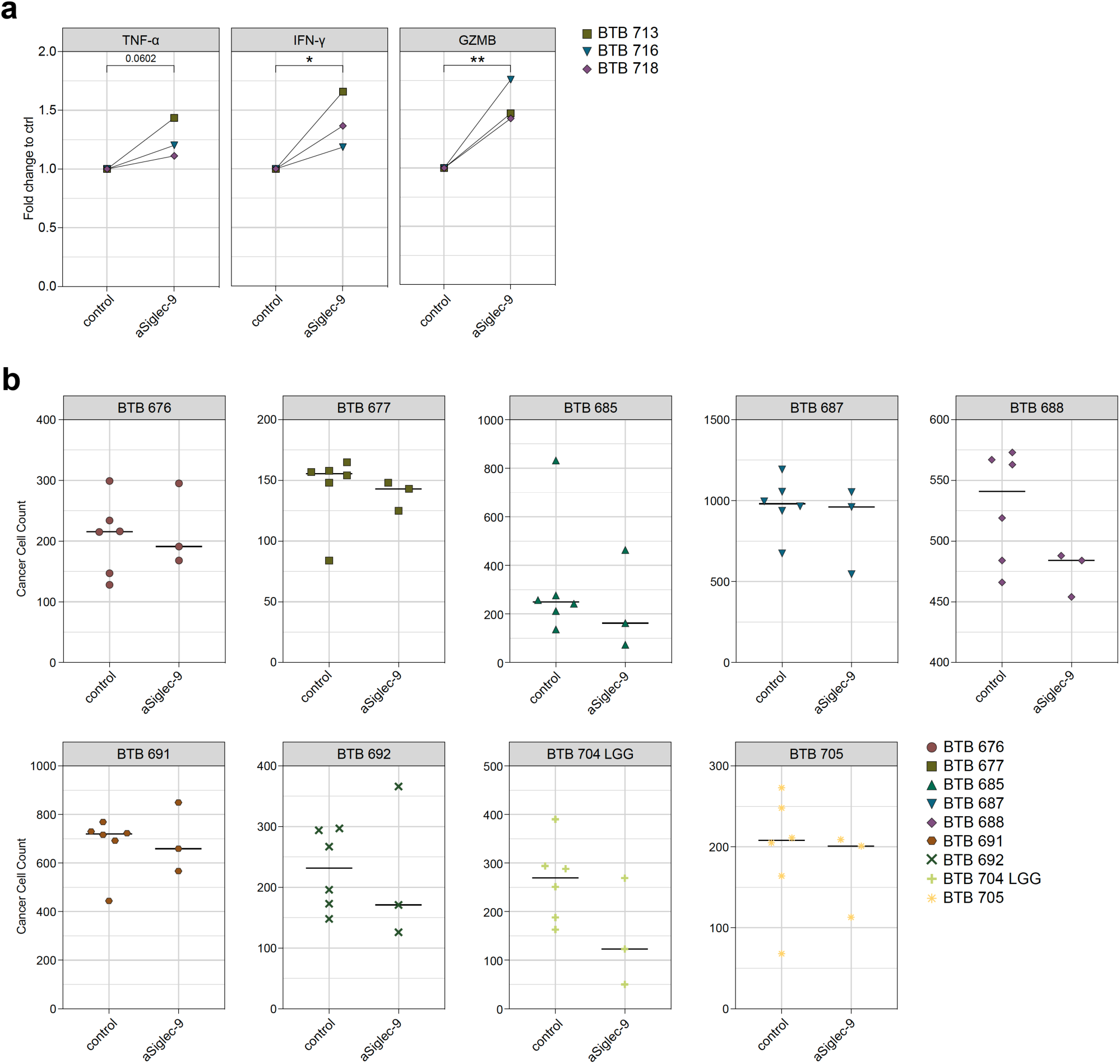
Siglec-9 blockade induces immune response and anti-tumor activity in human GBM explants. **a**, Fold change of TNF-α, IFN-γ and GZMB secretion measured in anti-Siglec-9 treated versus control bioreactor media in responding patients (response defined as fold change > 1 in all three comparisons). **b**, Number of glioma cells in the different treatment conditions for each patient obtained on a single-cell resolution from the image analyses. Individual symbols represent technical replicates within the treatment conditions. Statistics: Data are presented as median (**b**), unpaired two-tailed Student’s t test (**a**). *p ≤ 0.05, **p ≤ 0.01.

**Supplementary Data Fig. 7.**
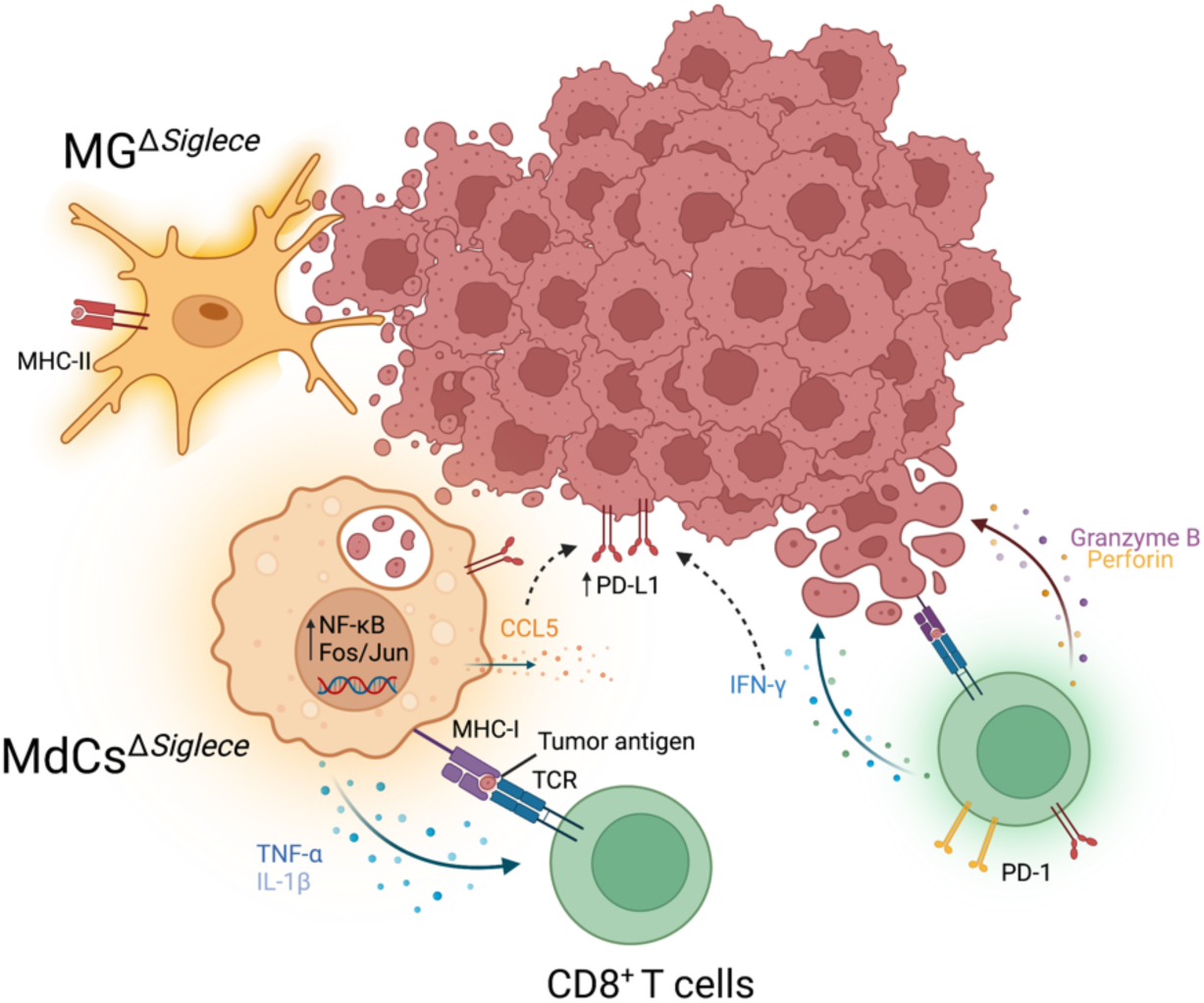
Summary of the findings described in this study.

