## Supplementary Table 1 for "The Siglec-sialic acid-axis is a target for innate immunotherapy of glioblastoma"

**Differentially expressed genes in cluster 15 MdC_Proliferating_Cre-pos vs. Cre-neg**

| GeneId | Symbol | GeneName | log2FC | log2A | P.Value | adj.P.Val |
|---|---|---|---|---|---|---|
| ENSMUSG00000062611 | Rps3a2 | ribosomal protein S3A2 | 3.256535368 | 7.222530341 | 1.14E-09 | 5.65E-06 |
| ENSMUSG00000059751 | Rps3a3 | ribosomal protein S3A3 | 2.902397786 | 5.749052657 | 2.86E-12 | 2.83E-08 |
| ENSMUSG00000091383 | Hist1h2al | histone cluster 1, H2al | 2.216514941 | 5.179374358 | 2.38E-08 | 5.75E-05 |
| ENSMUSG00000031880 | Rrad | Ras-related associated with diabetes | 1.370520205 | 5.022001293 | 1.53E-07 | 0.00025141 |
| ENSMUSG00000045193 | Cirbp | cold inducible RNA binding protein | 1.330264739 | 6.80543185 | 2.91E-08 | 5.75E-05 |
| ENSMUSG00000108366 | Gm5586 | predicted gene 5586 | 0.917072042 | 3.353926298 | 5.05E-06 | 0.006241098 |
| ENSMUSG00000036948 | Map11 | microtubule associated protein 11 | -1.327350384 | 6.456241487 | 4.56E-05 | 0.04505566 |
| ENSMUSG00000032501 | Trib1 | tribbles pseudokinase 1 | -1.331429568 | 7.281597405 | 1.99E-05 | 0.021882126 |
| ENSMUSG00000091993 | B930036N10Rik | RIKEN cDNA B930036N10 gene | -1.594833305 | 5.183668758 | 1.44E-06 | 0.002038023 |
| ENSMUSG00000095366 | Gm21860 | predicted gene, 21860 | -1.960480501 | 5.842608792 | 1.50E-08 | 4.93E-05 |

**Differentially expressed genes in cluster 16 TAMs_Cre-pos vs. Cre-neg**

| GeneId | Symbol | GeneName | log2FC | log2A | P.Value | adj.P.Val |
|---|---|---|---|---|---|---|
| ENSMUSG00000059751 | Rps3a3 | ribosomal protein S3A3 | 4.815379604 | 6.271008284 | 4.34E-07 | 0.001302366 |
| ENSMUSG00000062611 | Rps3a2 | ribosomal protein S3A2 | 4.276975217 | 7.842189586 | 3.58E-07 | 0.001302366 |
| ENSMUSG00000108366 | Gm5586 | predicted gene 5586 | 2.961356569 | 4.49630346 | 6.04E-06 | 0.00906393 |
| ENSMUSG00000045193 | Cirbp | cold inducible RNA binding protein | 1.371387985 | 6.677111821 | 1.16E-05 | 0.01158267 |
| ENSMUSG00000024042 | Sik1 | salt inducible kinase 1 | 0.688711953 | 6.635189368 | 5.50E-05 | 0.041240592 |
| ENSMUSG00000009013 | Dynll1 | dynein light chain LC8-type 1 | -0.71684861 | 9.229158046 | 3.22E-05 | 0.028996467 |
| ENSMUSG00000022957 | Itsn1 | intersectin 1 (SH3 domain protein 1A) | -0.72067051 | 5.88556613 | 8.89E-06 | 0.011430224 |
| ENSMUSG00000063317 | Usp31 | ubiquitin specific peptidase 31 | -0.959878739 | 4.071572574 | 4.22E-06 | 0.007589931 |
| ENSMUSG00000000489 | Pdgfb | platelet derived growth factor, B polypeptide | -1.04366702 | 6.211022667 | 1.95E-06 | 0.004389114 |
| ENSMUSG00000049539 | H1f1 | H1.1 linker histone, cluster member | -1.11191813 | 3.88917685 | 1.15E-05 | 0.01158267 |
| ENSMUSG00000069272 | H2ac8 | H2A clustered histone 8 | -1.147007282 | 6.188479021 | 2.45E-11 | 2.21E-07 |
| ENSMUSG00000020053 | Igf1 | insulin-like growth factor 1 | -1.869456214 | 4.615192321 | 4.38E-05 | 0.035849436 |

**Differentially expressed genes in cluster 17 MdC_Ly6c-low_CD11c-neg_Cre-pos vs. Cre-neg**

| GeneId | Symbol | GeneName | log2FC | log2A | P.Value | adj.P.Val |
|---|---|---|---|---|---|---|
| ENSMUSG00000062611 | Rps3a2 | ribosomal protein S3A2 | 3.893815201 | 7.852467057 | 3.02E-06 | 0.006386377 |
| ENSMUSG00000059751 | Rps3a3 | ribosomal protein S3A3 | 3.796629655 | 5.943324294 | 1.68E-07 | 0.00070972 |
| ENSMUSG00000108366 | Gm5586 | predicted gene 5586 | 2.531236748 | 4.824567236 | 1.72E-05 | 0.020072941 |
| ENSMUSG00000045193 | Cirbp | cold inducible RNA binding protein | 1.290701876 | 6.669848998 | 1.58E-06 | 0.004452299 |
| ENSMUSG00000097296 | Gm26532 | predicted gene, 26532 | 0.90062554 | 5.685734198 | 2.32E-05 | 0.020072941 |
| ENSMUSG00000031584 | Gsr | glutathione reductase | 0.813702917 | 7.257344292 | 1.90E-05 | 0.020072941 |
| ENSMUSG00000091017 | Fam71a | family with sequence similarity 71, member A | -1.004909372 | 5.130486638 | 1.54E-05 | 0.020072941 |
| ENSMUSG00000105954 | Gm42793 | predicted gene 42793 | -1.147247725 | 6.188398996 | 3.90E-09 | 3.30E-05 |
| ENSMUSG00000030117 | Gdf3 | growth differentiation factor 3 | -1.257976996 | 5.162756142 | 2.37E-05 | 0.020072941 |
| ENSMUSG00000050370 | Ch25h | cholesterol 25-hydroxylase | -1.31439953 | 4.624830363 | 1.27E-05 | 0.020072941 |

**Differentially expressed genes in cluster 18 Monocytes_Cre-pos vs. Cre-neg**

| GeneId | Symbol | GeneName | log2FC | log2A | P.Value | adj.P.Val |
|---|---|---|---|---|---|---|
| ENSMUSG00000062611 | Rps3a2 | ribosomal protein S3A2 | 3.622173482 | 7.614951928 | 9.11E-09 | 2.37E-05 |
| ENSMUSG00000059751 | Rps3a3 | ribosomal protein S3A3 | 3.513331908 | 5.702060408 | 3.97E-09 | 1.55E-05 |
| ENSMUSG00000108366 | Gm5586 | predicted gene 5586 | 2.438754048 | 4.676317849 | 6.43E-10 | 5.01E-06 |
| ENSMUSG00000027984 | Hadh | hydroxyacyl-Coenzyme A dehydrogenase | 0.922811888 | 5.097504703 | 1.22E-05 | 0.013279928 |
| ENSMUSG00000031709 | Tbc1d9 | TBC1 domain family, member 9 | 0.911333999 | 7.62894991 | 1.06E-06 | 0.001657745 |
| ENSMUSG00000047810 | Ccdc88b | coiled-coil domain containing 88B | -0.948029228 | 7.06864306 | 3.78E-06 | 0.004903711 |
| ENSMUSG00000066684 | Pilrb1 | paired immunoglobin-like type 2 receptor beta 1 | -0.950797639 | 5.479609185 | 3.17E-05 | 0.024720854 |
| ENSMUSG00000109498 | Gm45222 | predicted gene 45222 | -1.06083752 | 4.952431011 | 1.36E-05 | 0.013279928 |
| ENSMUSG00000095366 | Gm21860 | predicted gene, 21860 | -1.113400167 | 4.128894503 | 1.90E-05 | 0.01641656 |
| ENSMUSG00000094777 | Hist1h2ap | histone cluster 1, H2ap | -1.181846694 | 5.942599444 | 3.85E-05 | 0.027260106 |
| ENSMUSG00000028037 | Ifi44 | interferon-induced protein 44 | -1.305170267 | 5.204685793 | 7.32E-08 | 0.000142553 |
| ENSMUSG00000040026 | Saa3 | serum amyloid A 3 | -1.494577114 | 7.337815865 | 4.36E-05 | 0.028280697 |

**Differentially expressed genes in cluster 19 MdC_Ly6c-low_CD11c-pos_Cre-pos vs. Cre-neg**

| GeneId | Symbol | GeneName | log2FC | log2A | P.Value | adj.P.Val |
|---|---|---|---|---|---|---|
| ENSMUSG00000066553 | Gm6969 | predicted pseudogene 6969 | -2.809618532 | 5.37789483 | 1.70E-05 | 0.012472103 |
| ENSMUSG00000031494 | Cd209a | CD209a antigen | -1.771864621 | 6.345855703 | 7.39E-05 | 0.028135073 |
| ENSMUSG00000074896 | Ifit3 | interferon-induced protein with tetratricopeptide repeats 3 | -1.607046685 | 7.37183869 | 0.000257116 | 0.04581156 |
| ENSMUSG00000028037 | Ifi44 | interferon-induced protein 44 | -1.595600611 | 5.340962347 | 1.58E-05 | 0.012472103 |
| ENSMUSG00000035692 | Isg15 | ISG15 ubiquitin-like modifier | -1.537344472 | 9.559363648 | 1.53E-06 | 0.00256173 |
| ENSMUSG00000034459 | Ifit1 | interferon-induced protein with tetratricopeptide repeats 1 | -1.406962827 | 6.215815838 | 0.000217245 | 0.042150581 |
| ENSMUSG00000073491 | Ifi213 | interferon activated gene 213 | -1.356120554 | 7.186425438 | 6.80E-06 | 0.008116903 |
| ENSMUSG00000045932 | Ifit2 | interferon-induced protein with tetratricopeptide repeats 2 | -1.333143362 | 7.372967778 | 0.000151808 | 0.033016729 |
| ENSMUSG00000022584 | Ly6c2 | lymphocyte antigen 6 complex, locus C2 | -1.314348572 | 9.313851975 | 6.15E-05 | 0.026683467 |
| ENSMUSG00000079017 | Ifi27l2a | interferon, alpha-inducible protein 27 like 2A | -1.280867136 | 10.00107306 | 2.12E-05 | 0.014484054 |
| ENSMUSG00000030107 | Usp18 | ubiquitin specific peptidase 18 | -1.256248733 | 6.740330743 | 2.71E-05 | 0.017285854 |
| ENSMUSG00000039236 | Isg20 | interferon-stimulated protein | -1.242277602 | 7.426720509 | 1.33E-07 | 0.000424975 |
| ENSMUSG00000052749 | Trim30b | tripartite motif-containing 30B | -1.167563734 | 4.896874117 | 4.60E-05 | 0.022431098 |
| ENSMUSG00000032690 | Oas2 | 2'-5' oligoadenylate synthetase 2 | -1.135499859 | 5.867241537 | 1.57E-05 | 0.012472103 |
| ENSMUSG00000024675 | Ms4a4c | membrane-spanning 4-domains, subfamily A, member 4C | -1.131510771 | 8.934039529 | 0.000171349 | 0.035584747 |
| ENSMUSG00000057596 | Trim30d | tripartite motif-containing 30D | -1.115597345 | 5.977623542 | 0.00013132 | 0.031362389 |
| ENSMUSG00000052776 | Oas1a | 2'-5' oligoadenylate synthetase 1A | -1.112954894 | 7.623783764 | 0.000109058 | 0.029396481 |
| ENSMUSG00000073489 | Ifi204 | interferon activated gene 204 | -1.107071873 | 7.125669016 | 0.000284229 | 0.048395536 |
| ENSMUSG00000106734 | Gm20559 | predicted gene, 20559 | -1.048168826 | 4.35965969 | 0.000159152 | 0.033786167 |
| ENSMUSG00000040296 | Ddx58 | DEAD (Asp-Glu-Ala-Asp) box polypeptide 58 | -1.027664122 | 6.502970863 | 0.000288762 | 0.048395536 |
| ENSMUSG00000026536 | Ifi211 | interferon activated gene 211 | -0.999414608 | 8.087154466 | 0.000220614 | 0.042150581 |
| ENSMUSG00000091649 | Phf11b | PHD finger protein 11B | -0.994686201 | 8.001115743 | 8.15E-05 | 0.028716677 |
| ENSMUSG00000043099 | Hic1 | hypermethylated in cancer 1 | -0.988425655 | 5.6567225 | 6.90E-05 | 0.028135073 |
| ENSMUSG00000019122 | Ccl9 | chemokine (C-C motif) ligand 9 | -0.967135597 | 6.134711267 | 7.12E-05 | 0.028135073 |
| ENSMUSG00000027514 | Zbp1 | Z-DNA binding protein 1 | -0.918070038 | 7.542218137 | 8.42E-05 | 0.028716677 |
| ENSMUSG00000027078 | Ube2l6 | ubiquitin-conjugating enzyme E2L 6 | -0.909910982 | 6.962280526 | 0.000110779 | 0.029396481 |
| ENSMUSG00000021453 | Gadd45g | growth arrest and DNA-damage-inducible 45 gamma | -0.905902445 | 7.688525527 | 0.00014121 | 0.032226927 |
| ENSMUSG00000046718 | Bst2 | bone marrow stromal cell antigen 2 | -0.900029377 | 9.930186082 | 7.66E-05 | 0.028135073 |
| ENSMUSG00000075269 | Bex6 | brain expressed family member 6 | -0.882179203 | 3.799169313 | 0.000127066 | 0.031124692 |

| | | | | | | |
|---|---|---|---|---|---|---|
| ENSMUSG00000033355 | Rtp4 | receptor transporter protein 4 | -0.881121415 | 7.135211259 | 0.000109875 | 0.029396481 |
| ENSMUSG00000036986 | Pml | promyelocytic leukemia | -0.873275229 | 6.388467558 | 0.000277927 | 0.04827342 |
| ENSMUSG00000091017 | Fam71a | family with sequence similarity 71, member A | -0.855121212 | 4.644112139 | 5.13E-05 | 0.023326072 |
| ENSMUSG00000015947 | Fcgr1 | Fc receptor, IgG, high affinity I | -0.789370086 | 7.552478095 | 1.19E-05 | 0.011352186 |
| ENSMUSG00000070284 | Gmppb | GDP-mannose pyrophosphorylase B | -0.680418978 | 5.699560694 | 0.000105784 | 0.029396481 |
| ENSMUSG00000020277 | Pfkl | phosphofructokinase, liver, B-type | -0.665323261 | 6.743177969 | 0.000199696 | 0.040050591 |
| ENSMUSG00000026946 | Nmi | N-myc (and STAT) interactor | -0.66010306 | 6.645891092 | 0.00012035 | 0.031073093 |
| ENSMUSG00000009013 | Dynll1 | dynein light chain LC8-type 1 | -0.658962438 | 8.980017225 | 0.000152071 | 0.033016729 |
| ENSMUSG00000026979 | Psd4 | pleckstrin and Sec7 domain containing 4 | -0.633314698 | 6.102406472 | 0.000107748 | 0.029396481 |
| ENSMUSG00000020841 | Cpd | carboxypeptidase D | 0.700037101 | 6.451338697 | 0.000141686 | 0.032226927 |
| ENSMUSG00000024248 | Cox7a2l | cytochrome c oxidase subunit 7A2 like | 0.749404698 | 8.760440964 | 0.000110234 | 0.029396481 |
| ENSMUSG00000060147 | Serpinb6a | serine (or cysteine) peptidase inhibitor, clade B, member 6a | 0.784469369 | 4.724863605 | 3.16E-05 | 0.018878258 |
| ENSMUSG00000040680 | Kremen2 | kringle containing transmembrane protein 2 | 0.804923538 | 3.714157136 | 0.000258958 | 0.04581156 |
| ENSMUSG00000058997 | Vwa8 | von Willebrand factor A domain containing 8 | 0.909679621 | 4.888839904 | 2.81E-06 | 0.003836597 |
| ENSMUSG00000058216 | Gstp3 | glutathione S-transferase pi 3 | 0.952587507 | 4.042536854 | 4.43E-05 | 0.022431098 |
| ENSMUSG00000061175 | Fnip2 | folliculin interacting protein 2 | 1.013308623 | 3.985362874 | 0.000231666 | 0.043394143 |
| ENSMUSG00000026070 | Il18r1 | interleukin 18 receptor 1 | 1.026247236 | 3.775059653 | 1.03E-05 | 0.010907821 |
| ENSMUSG00000030532 | Hddc3 | HD domain containing 3 | 1.11374411 | 3.635163066 | 9.78E-05 | 0.029396481 |
| ENSMUSG00000045193 | Cirbp | cold inducible RNA binding protein | 1.121527944 | 7.474437834 | 4.70E-05 | 0.022431098 |
| ENSMUSG00000051517 | Arhgef39 | Rho guanine nucleotide exchange factor (GEF) 39 | 1.183256865 | 3.940181889 | 0.000110151 | 0.029396481 |
| ENSMUSG00000037940 | Inpp4b | inositol polyphosphate-4-phosphatase, type II | 1.376140536 | 4.566312095 | 1.61E-06 | 0.00256173 |
| ENSMUSG00000024910 | Ctsw | cathepsin W | 1.420815969 | 4.724172323 | 3.16E-08 | 0.000302291 |
| ENSMUSG00000002033 | Cd3g | CD3 antigen, gamma polypeptide | 1.476588812 | 4.938332045 | 0.000126767 | 0.031124692 |
| ENSMUSG00000032093 | Cd3e | CD3 antigen, epsilon polypeptide | 1.760479238 | 4.775774843 | 0.000248792 | 0.045705935 |
| ENSMUSG00000091383 | Hist1h2al | histone cluster 1, H2al | 2.304015545 | 4.22635166 | 0.000201238 | 0.040050591 |
| ENSMUSG00000108366 | Gm5586 | predicted gene 5586 | 2.624449935 | 4.92628155 | 4.45E-05 | 0.022431098 |
| ENSMUSG00000062611 | Rps3a2 | ribosomal protein S3A2 | 4.212674479 | 8.593788693 | 1.22E-07 | 0.000424975 |
| ENSMUSG00000059751 | Rps3a3 | ribosomal protein S3A3 | 4.225108755 | 6.085079729 | 5.46E-07 | 0.001302822 |
