## Supplementary Table 2-4 for "The Siglec-sialic acid-axis is a target for innate immunotherapy of glioblastoma"

**Supplementary Table 2.** Patient characteristics

| Patient | Sex | Age | Histology | Grade | Status | IDH | MGMT promotor | EGFR | Subclass | Study |
|---|---|---|---|---|---|---|---|---|---|---|
| BTB 700R | M | 51 | GBM | IV | recurrent | WT | methylated | amplified | classical | Bioreactor |
| BTB 713 | F | 47 | GBM | IV | primary | WT | methylated | not amplified | mesenchymal | Bioreactor |
| BTB 714 | M | 61 | GBM | IV | primary | WT | unmethylated | not amplified | proneural | Bioreactor |
| BTB 716 | M | 72 | GBM | IV | primary | WT | unmethylated | not amplified | proneural | Bioreactor |
| BTB 718 | M | 59 | GBM | IV | primary | WT | methylated | amplified | classical | Bioreactor |
| BTB 676 | M | 71 | GBM | IV | primary | WT | methylated | amplified | classical | Pharmacos |
| BTB 677 | M | 58 | GBM | IV | primary | WT | unmethylated | not amplified | classical | Pharmacos |
| BTB 685 | M | 63 | GBM | IV | primary | WT | unmethylated | not amplified | proneural | Pharmacos |
| BTB 687 | F | 54 | GBM | IV | primary | WT | methylated | amplified | proneural | Pharmacos |
| BTB 688 | F | 72 | GBM | IV | primary | WT | methylated | not amplified | mesenchymal | Pharmacos |
| BTB 691 | M | 63 | GBM | IV | primary | WT | unmethylated | not amplified | proneural | Pharmacos |
| BTB 692 | M | 70 | GBM | IV | primary | WT | methylated | amplified | classical | Pharmacos |
| BTB 704 | M | 62 | LGG | I-II | primary | WT | unmethylated | not amplified | LGG | Pharmacos |
| BTB 705 | F | 78 | GBM | IV | primary | WT | methylated | amplified | classical | Pharmacos |

**Supplementary Table 3.** Anti-mouse Flow Cytometry Antibodies

| Specificity | Clone | Fluorochrome | Manufacturer | Catalog # |
|---|---|---|---|---|
| CD45 | 30-F11 | FITC | BioLegend | 103107 |
| CD45 | 30-F11 | BV605 | BioLegend | 103140 |
| CD45 | 30-F11 | BV421 | BioLegend | 103134 |
| CD45 | 30-F11 | PE | BioLegend | 103106 |
| CD45.1 | A20 | PE-Cy7 | BioLegend | 110730 |
| CD11b | M1/70 | BV650 | BioLegend | 101259 |
| CD11b | M1/70 | APC-Cy7 | BioLegend | 101226 |
| Cx3cr1 | SA011F11 | FITC | BioLegend | 149020 |
| Siglec-E | M1304A01 | APC | BioLegend | 677106 |
| Siglec-E | M1304A01 | PE | BioLegend | 677104 |
| Siglec-E | M1304A01 | PerCP-Cy5.5 | BioLegend | 677114 |
| CD163 | S15049l | BV421 | BioLegend | 155309 |
| CD163 | S15049l | PE-Cy7 | BioLegend | 155320 |
| CD86 | GL-1 | APC-Cy7 | BioLegend | 105030 |
| CD80 | 16-10A1 | PE-Cy7 | BioLegend | 104734 |
| CD80 | 16-10A1 | BV650 | BioLegend | 104732 |
| I-A/I-E (MHC class II) | M5/114.15.2 | PerCP-Cy5.5 | BioLegend | 107626 |
| NK-1.1 | PK136 | PE-Cy5 | BioLegend | 108715 |
| Ly-6G/Ly-6C (Gr-1) | RB6-8C5 | AF647 | BioLegend | 108418 |
| CD3e | 145-2C11 | PE-Cy7 | BioLegend | 100319 |
| CD3e | 145-2C11 | APC-Cy7 | BioLegend | 100330 |
| CD4 | RM4-5 | PerCP-Cy5.5 | BioLegend | 100540 |
| CD8a | 53-6.7 | FITC | BioLegend | 100706 |
| CD8a | 53-6.7 | PE-Cy7 | BioLegend | 100722 |
| CD8a | 53-6.7 | PerCP-Cy5.5 | BioLegend | 100734 |
| PD-1 | 29F.1A12 | APC-Cy7 | BioLegend | 135224 |
| TIM-3 | RMT3-23 | APC | BioLegend | 119705 |
| LAG-3 | C9B7W | BV421 | BioLegend | 125221 |
| CTLA-4 | UC10-4B9 | PE | BioLegend | 106306 |
| PD-L1 | 10F.9G2 | BV650 | BioLegend | 124336 |
| IFN-γ | XMG 1.2 | FITC | BioLegend | 505806 |
| IFN-γ | XMG 1.2 | APC-Cy7 | BioLegend | 505850 |
| IFN-γ | XMG 1.2 | PE | BioLegend | 505808 |
| TNF-α | MP6-XT22 | APC | BioLegend | 506308 |
| Ki-67 | 16A8 | APC | BioLegend | 652406 |
| Ki-67 | 16A8 | PE-Cy7 | BioLegend | 652426 |
| CD25 | PC61 | APC | BioLegend | 102012 |
| CD69 | H1.2F3 | BV650 | BioLegend | 104541 |
| CD69 | H1.2F3 | PerCP-Cy5.5 | BioLegend | 104522 |

| | | | | |
|---|---|---|---|---|
| CD44 | IM7 | PE | BioLegend | 103008 |
| CD107a | 1D4B | PE-Cy7 | BioLegend | 121620 |
| Cxcr3 | CXCR3-173 | BV650 | BioLegend | 126531 |
| Foxp3 | MF-14 | AF488 | BioLegend | 126406 |
| Foxp3 | MF-14 | PE | BioLegend | 126404 |
| T-bet | 4B10 | PE-Cy7 | BioLegend | 644824 |
| Siglec-H | 551 | PerCP-Cy5.5 | BioLegend | 129614 |
| CD22 | OX-97 | FITC | BioLegend | 126106 |
| Siglec-G | SH2.1 | APC | eBioscience | 17-5833-82 |
| H-2Kb | AF6-88.5 | APC | BioLegend | 116518 |
| H-2Kb-SIINFEKL | 25-D1.16 | APC | eBioscience | 17-5743-82 |
| IgG2a | RMG2a-62 | APC | BioLegend | 407110 |
| IgG2a | RMG2a-62 | BV421 | BioLegend | 407117 |
| IgG | Polyclonal | AF488 | Invitrogen | A32723 |

**Supplementary Table 4.** Anti-human Flow Cytometry Antibodies

| Specificity | Clone | Fluorochrome | Manufacturer | Catalog # |
|---|---|---|---|---|
| CD45 | 2D1 | BV510 | BioLegend | 368526 |
| CD11b | ICRF44 | BV650 | BioLegend | 301336 |
| CX3CR1 | 2A9-1 | PE-Cy7 | BioLegend | 341612 |
| TMEM119 | A16075D | - | BioLegend | 853302 |
| Siglec-1 | 7-239 | AF647 | BioLegend | 346006 |
| CD22 | HIB22 | AF647 | BioLegend | 302518 |
| CD33 | HIM3-4 | FITC | BioLegend | 303304 |
| Siglec-5 | 1A5 | PE | BioLegend | 352004 |
| Siglec-7 | 6-434 | PE | BioLegend | 339204 |
| Siglec-8 | 7C9 | APC | BioLegend | 347106 |
| Siglec-9 | K8 | APC | BioLegend | 351506 |
| Siglec-10 | 5G6 | PE | BioLegend | 347603 |
